# Phage-based microbiome manipulation reveals ecological interactions within gut communities

**DOI:** 10.64898/2026.05.13.724931

**Authors:** Taylor H. Nguyen, Morgan Su, Nhien T. Lu, Valentine Trotter, Saria A. McKeithen-Mead, Jamie Alcira Lopez, Jiawei Sun, Zachary Hallberg, Handuo Shi, Po-Yi Ho, Brian C. DeFelice, Michiko E. Taga, Adam M. Deutschbauer, Andrew J. Hryckowian, Kerwyn Casey Huang

## Abstract

Mechanistic understanding of gut ecology is limited by the availability of tools for precise manipulation of microbiome composition. Here, we isolate lytic phages to enable targeted removal of gut commensal *Escherichia fergusonii* (*Ef*) from complex, undefined stool-derived *in vitro* communities. A single phage drove resistance without fitness cost in monoculture, but resistant *Ef* exhibited reduced fitness in communities, enabling expansion of closely related Proteobacteria. Resistance arose via reversible promoter inversion linked to outer-membrane function. A phage cocktail overcame resistance to achieve *Ef* knockout across communities with minimal collateral effects. Using knockout communities, we show that *Ef* is necessary and sufficient for preventing *Salmonella* invasion. Replacement with an *Ef* transposon-mutant library revealed that community-specific fitness defects are enriched in genes involved in outer-membrane assembly. Disruption of these genes sensitized *Ef* to antagonistic community members, highlighting interspecies warfare as a key driver of microbiome ecology. These results establish phage-mediated perturbation as a framework for linking species to community-level function and for enabling precision microbiome engineering.

## Introduction

As associations between gut microbiome composition and various disease states^1,2^ have become increasingly clear, efforts to engineer gut microbial communities as therapeutic interventions have accelerated. Environmental stresses^3–6^, diet^7–9^, and chemical perturbations^10,11^ can alter microbiome composition, but these approaches are typically broad and difficult to control, limiting their utility for targeted manipulation. Antibiotics, in particular, can cause long-lasting collateral damage to the microbiome^5^ with adverse consequences for host health^12,13^, while the rise of antibiotic resistance poses a growing global threat^14,15^. Despite the development of alternative modalities, including antimicrobial peptides^16,17^, small molecules^18^, and antibodies^19^, tools for precise, species-level manipulation of complex microbial communities remain lacking. New strategies are therefore needed to enable targeted and predictable microbiome engineering^20–22^.

Lytic bacteriophages (“phages”), viruses that infect and lyse bacteria, offer a promising route to such precision due to their high specificity for a species, strain, or even a particular phenotypic variant of a strain^23–25^. Although phage therapy predates antibiotics and has long been used to treat infections^26,27^, its broader potential has remained underexplored, particularly in complex microbial communities. Phages are abundant^28^ and are readily detected in the gut microbiome by metagenomic sequencing^29^, yet their ecological impacts in these environments remain poorly understood^30^. While recent efforts have expanded collections of gut phages^23,24,31^, lytic phages have not been isolated for most commensal species, and it remains unclear whether phages can be harnessed to manipulate microbiome composition in a predictable and controllable manner.

Beyond their therapeutic potential^32–34^, phages provide a unique opportunity to probe ecological interactions within microbial communities. Most studies of phage–bacteria dynamics have been performed in monoculture, yet emerging evidence indicates that community context strongly influences the evolution of phage resistance^35^ and its consequences^36^. Moreover, phage treatment can restructure communities indirectly, even when non-target species are not directly susceptible^37^, highlighting the complexity and potential unpredictability of phage-driven ecological dynamics.

These challenges motivate the development of approaches that combine the specificity of phages with experimental systems that preserve community complexity. *In vitro* communities have emerged as powerful models of the gut microbiome, enabling interrogation of ecological interactions that drive community functions, including responses to drugs^10,11^ and colonization resistance to pathogens^38,39^. Top-down, stool-derived communities retain much of the diversity and ecological structure of native microbiotas *in vivo*^38,40^ but are difficult to interrogate mechanistically due to their complexity. In contrast, bottom-up synthetic communities offer greater experimental control but are typically comprised of a limited set of abundant, representative species and therefore lack strain diversity and low-abundance members that may play critical roles in community function^41–43^. Phage-based removal of individual species provides a complementary strategy: by generating targeted “knockout” communities within complex backgrounds, it becomes possible to directly test the roles of specific species and genes in community assembly, microbial interactions, and community-level functions. More broadly, approaches enable causal interrogation of microbiome ecology across molecular, genetic, and community scales.

Here, we establish phage-based species knockout as a general strategy to dissect microbial community function. Using stool-derived *in vitro* communities, we isolate lytic phages that target the commensal *Escherichia fergusonii* (*Ef*) and show that resistance arises rapidly but incurs a fitness cost specifically in communities, enabling other phylogenetically similar species to expand and reshaping competitive dynamics. We identify a reversible promoter inversion upstream of the *hyx* operon as a key resistance mechanism and show that phage cocktails can overcome resistance to achieve robust *Ef* knockout with minimal disruption to overall community composition. Leveraging these knockout communities, we demonstrate that *Ef* is both necessary and sufficient for colonization resistance against *Salmonella* and use genome-scale transposon mutagenesis to uncover outer membrane assembly as a critical determinant of fitness in community contexts. Together, these results link phage resistance, envelope remodeling, and community-dependent fitness, revealing the presence of ecological warfare within gut communities and establishing phage-based editing as a powerful framework for connecting species and genes to ecological function in complex microbiomes.

## Results

### Phage resistance arises rapidly without fitness cost in monoculture

To establish phage-based manipulation of microbial communities, we targeted the abundant Enterobacteriaceae member *Ef* in stool-derived communities that we previously established as an experimental model system for the gut microbiome^11,38^. These communities preserve family-level abundance patterns observed *in vivo*^38^. One notable exception is *Ef*, which is typically at low abundance *in vivo* but blooms *in vitro* to become dominant, motivating its removal to better model native community structure. Because Enterobacteriaceae blooms are associated with dysbiosis^44^, this system enables interrogation of the ecological consequences of targeting an abundant species. Using plaque assays (**Methods**), we isolated a lytic phage (Tay4; **Fig. 1a**, **Table S1**) targeting an isolate of the sole *Ef* strain detected in these communities^41^. Sequencing revealed that Tay4 belongs to the Caudoviricetes class of tailed lytic phages and is classified within the *Phapecoctavirus* genus (**Methods**).

**Figure 1:**
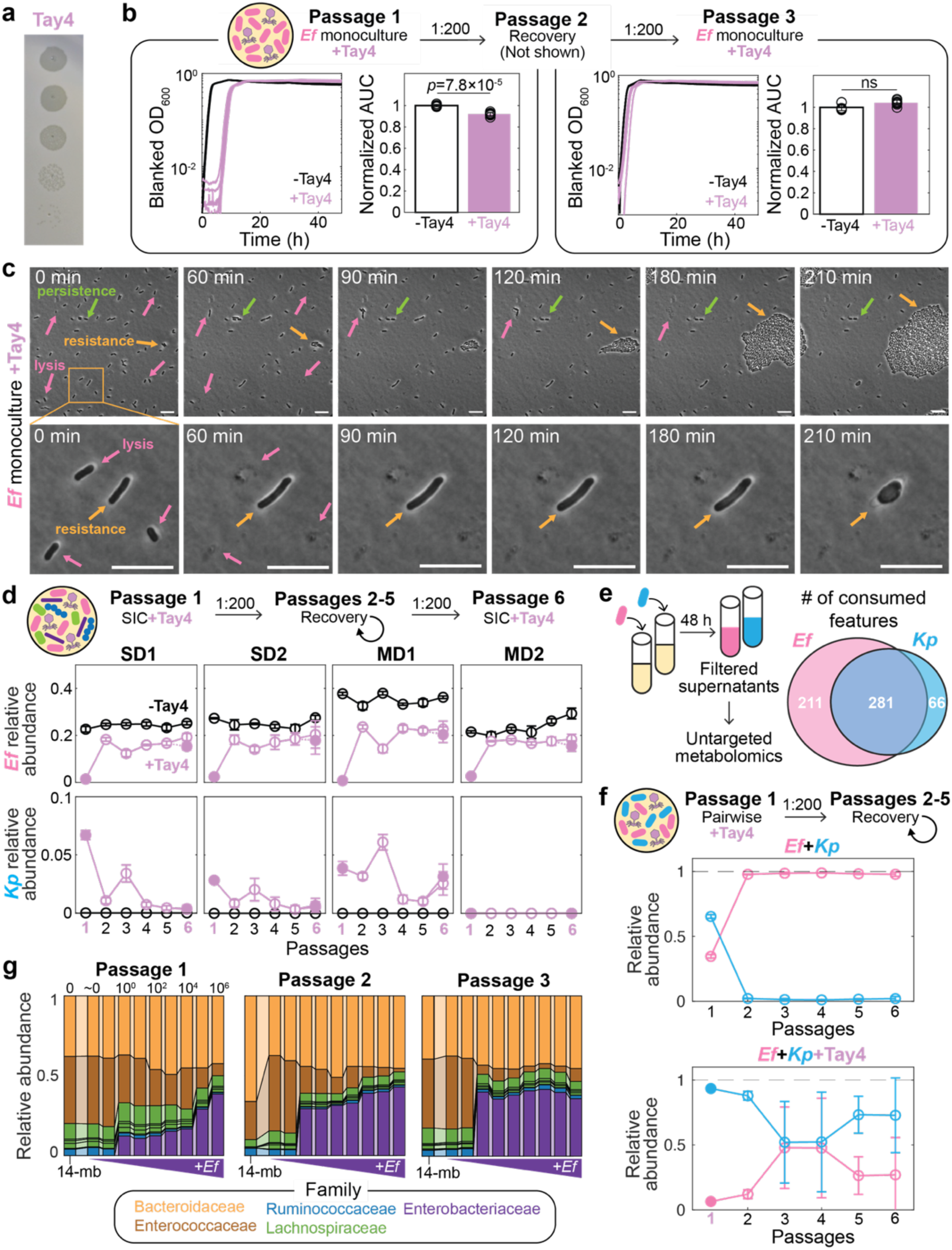
Phage-resistant *Escherichia fergusonii* exhibits a fitness defect specific to growth in microbial communities. a) Plaques of the lytic phage Tay4 on an *E. fergusonii* (*Ef*) lawn. Ten-fold serial dilutions of Tay4 (starting at 10^6^ PFUs) were spotted onto a BHI soft agar (0.35%) overlay on BHI agar (1.5%). b) Growth dynamics for *Ef* with (purple) and without (black) Tay4 (MOI∼1). Cultures were grown in passage 1, passaged 1:200 into phage-free medium for recovery (passage 2), and re-exposed to Tay4 (MOI∼1) in passage 3. Curves represent biological replicates (*n*=8). Area under the curve (AUC; 0–48 h) was normalized to the mean AUC of untreated *Ef*. *p*-values were calculated using a one-sided Wilcoxon rank-sum test (ns: not significant, *p* > 0.05). c) Time-lapse microscopy of *Ef* grown anaerobically with Tay4 (MOI∼1). Arrows indicate representative phenotypes: lysis (pink), persistence-like behavior (green), and resistance-associated growth or morphological changes (orange; zoomed-in at bottom). See **Movie S1** for full time-lapse. Scale bar: 10 µm. d) Relative abundance dynamics of *Ef* (top) and *Kp* (bottom) across stool-derived *in vitro* communities (SICs). Communities were treated with Tay4 (purple, MOI∼1) or left untreated (black) in passage 1, followed by four passages without the addition of phage for recovery. In passage 6, previously treated communities were re-exposed to Tay4 (dotted lines) or left untreated (solid lines) to assess resistance. Filled purple circles indicate passages with phage treatment. Points show mean±standard deviation (s.d.; *n*=3 biological replicates). e) Overlap in metabolite consumption between *Ef* and *Kp*. Venn diagram shows features uniquely or jointly depleted during monoculture growth. Features were defined as consumed if they decreased >100-fold relative to fresh BHI, had peak area >5,000 in BHI, and were significantly depleted (two-sample Student’s *t*-test with Benjamini-Hochberg correction, adjusted *p*<0.05). f) Pairwise competition between *Ef* (pink) and *Kp* (blue). Cultures were inoculated at a 1:10 *Ef*:*Kp* ratio and grown without (top) or with (bottom) Tay4 (MOI∼1) in passage 1, followed by four passages without added phage. Points show mean±s.d. (*n*=3 biological replicates). g) Steady-state composition of a 15-member synthetic community is insensitive to initial *Ef* abundance. Communities lacking *Ef* (14-member baseline; 14-mb) were supplemented with *Ef* across a range of input abundances (∼10^0^–10^6^ CFUs) and passaged two times (1:200 dilution). Bars show relative abundance of ASVs, colored by family.

When treated with Tay4 at a multiplicity of infection (MOI) ∼1, *Ef* growth was inhibited for ∼6 h due to phage-mediated lysis (**Fig. 1b,c**, **Fig. S1a**, **Movie S1**), as heat-denatured phage had no effect (**Fig. S1b**). Following this delay, cultures recovered and grew with kinetics similar to untreated controls, reflecting expansion of a resistant subpopulation (**Fig. S1a**, **Movie S1**), with additional single-cell behaviors consistent with persistence^45^ and envelope remodeling^46^ (**Fig. 1c**, **Movie S1**). We confirmed that Tay4 could not create plaques on a lawn of *Ef* after one passage of phage treatment (**Fig. S1c)**. Hereafter, to capture *Ef*’s dynamic response to phage treatment, we performed passages by diluting saturated cultures 1:200 into fresh medium and allowing them to grow for 48 h. After passaging in the presence of phage, these cultures retained resistance and exhibited growth dynamics comparable to untreated *Ef* (**Fig. S1d**), indicating that a substantial fraction of surviving cells had heritable resistance. When passaged once without phage and then re-exposed, cultures showed only a modest increase in lag time relative to untreated controls (**Fig. 1b**), consistent with partial maintenance of resistance in the absence of selection. Notably, phage-resistant *Ef* exhibited no detectable fitness cost in monoculture, as measured by growth rate, maximum optical density (OD), and biomass yield (**Fig. 1b, S1a**), indicating no detectable fitness cost in monoculture.

### Phage resistance incurs a fitness cost in communities, reshaping interspecies competition

We next asked whether phage resistance has different consequences in a community context. We hypothesized that the initial growth inhibition of *Ef* by phage treatment, combined with ecological competition, could enable out-competition of resistant subpopulations^36^. To test this hypothesis, we applied Tay4 to four stool-derived *in vitro* communities (SICs) from humanized mice, which originate from the same fecal inoculum but differ in composition due to host diet (standard diet (SD) and microbiota-accessible carbohydrate-deficient diet (MD); SD1, SD2, MD1, MD2)^38^. No Enterobacteriaceae other than *Ef* were detectable (relative abundance >10^-3^) in these SICs. Consistent with its specificity, Tay4 did not inhibit the monoculture growth of 15 other community isolates (**Fig. S2**). We treated each SIC with Tay4 and tracked community composition across treatment and recovery passages, followed by re-exposure to phage to assess resistance. Community biomass was similar with and without phage treatment (**Fig. S3a,b**), indicating that relative abundances are reasonable proxies for absolute abundances^41^. Following treatment, *Ef* abundance decreased sharply but subsequently recovered, indicating the emergence of heritable resistance (**Fig. 1d**). However, in contrast to monoculture (**Fig. 1b**), *Ef* consistently stabilized at a lower abundance across all communities (**Fig. 1d**), revealing that resistance incurs a fitness cost in community settings.

The primary compositional change following phage treatment was the emergence of another Enterobacteriaceae species, *Klebsiella pneumoniae* (*Kp*), which rose from below the limit of detection to ∼1% abundance in three of four communities and persisted after recovery of resistant *Ef* (**Fig. 1d**, **S3c**). This expansion suggests that resistance-associated fitness costs in *Ef* open ecological niches that can be occupied by closely related competitors. Consistent with this interpretation, metabolomics analysis^47^ revealed substantial niche overlap between *Ef* and *Kp*, with a subset of metabolites uniquely consumed by *Kp* (**Fig. 1e**), supporting its persistence at low abundance in the presence of wild-type *Ef*. In the community in which *Kp* did not emerge (MD2), *Ef* nonetheless stabilized at reduced abundance, indicating that the fitness defect is intrinsic to resistance rather than dependent on *Kp*, which instead expands into niches opened by this fitness cost. No other taxa exhibited comparable expansion (**Fig. S3c**), indicating that phage-mediated trade-offs selectively reshape competition within the Enterobacteriaceae.

To directly test the impact of phage resistance on interspecies competition, we co-cultured *Ef* and *Kp* with and without Tay4 treatment. In the absence of phage, *Ef* rapidly dominated the co-culture despite starting at 10-fold lower abundance (**Fig. 1f**), consistent with its competitive advantage. In contrast, phage treatment reversed this outcome, with *Kp* increasing in relative abundance following the initial depletion of *Ef* (**Fig. 1f**). These results demonstrate that resistance-associated fitness costs in *Ef* are sufficient to alter competitive outcomes, enabling expansion of closely related competitors.

We next considered the alternative possibility that the reduced abundance of *Ef* in communities reflects a transient effect of initial phage-mediated depletion rather than a true fitness cost. To test how quickly *Ef* can recover to its steady-state abundance from low starting levels, we used a synthetic 15-member community isolated from the SICs that recapitulates the composition of MD2^41^. We introduced *Ef* into the corresponding stabilized 14-member community (**Methods**) across a range of initial abundances and passaged the communities three times. Although *Ef* relative abundance after the first passage was higher for larger inocula, by the third passage it converged to ∼40% even when introduced at very low levels (starting inoculum of <10 cells; **Fig. 1g**). Only the most extreme dilutions failed to recover *Ef*, consistent with inocula that likely lacked *Ef* cells altogether because of bottlenecking.

We then repeated this experiment with concurrent addition of Tay4 and *Ef*. After three passages, *Ef* stabilized at ∼50% of its steady-state abundance in untreated communities (**Fig. S3d**), indicating a persistent fitness defect due to phage resistance. This effect was comparable to that observed in the undefined SICs (**Fig. 1d**) and occurred in the absence of *Kp*, consistent with the MD2 community. Together, these results indicate that *Ef* can rapidly regain its steady-state abundance after strong depletion, arguing that the reduced abundance of Tay4-treated *Ef* in communities is unlikely to reflect slow recovery alone. Instead, phage resistance imposes a stable fitness trade-off that can modulate community composition even without complete species removal.

### Phage resistance is reversible and growth-phase dependent

To interrogate why phage resistance confers a fitness disadvantage in communities, we sought to isolate Tay4-resistant *Ef* clones. We treated an *Ef* monoculture and the MD2 community with Tay4 and plated for single colonies (**Fig. S4a,b**; **Methods**). Although population-level dynamics indicated resistance (**Fig. 1b**, **Fig. 1d**, **S1a**), all isolated colonies exhibited a ∼6 h growth delay upon re-exposure to Tay4 (**Fig. S4a,b**), similar to wild-type *Ef* (**Fig. 1b**), indicating re-sensitization. These results suggest that resistance to Tay4 is not stably maintained during colony isolation.

We therefore hypothesized that Tay4 coexists with *Ef* through reversible transitions between resistant and susceptible states^48^. To test for coexistence, we quantified phage abundance during passaging of resistant cultures. Following initial infection, Tay4 titers increased ∼100-fold and remained high across multiple passages without exogenous phage addition in both monoculture and community settings (**Fig. 2a**). This persistence is inconsistent with a fully resistant population and instead indicates ongoing phage replication, likely sustained by a susceptible subpopulation.

**Figure 2:**
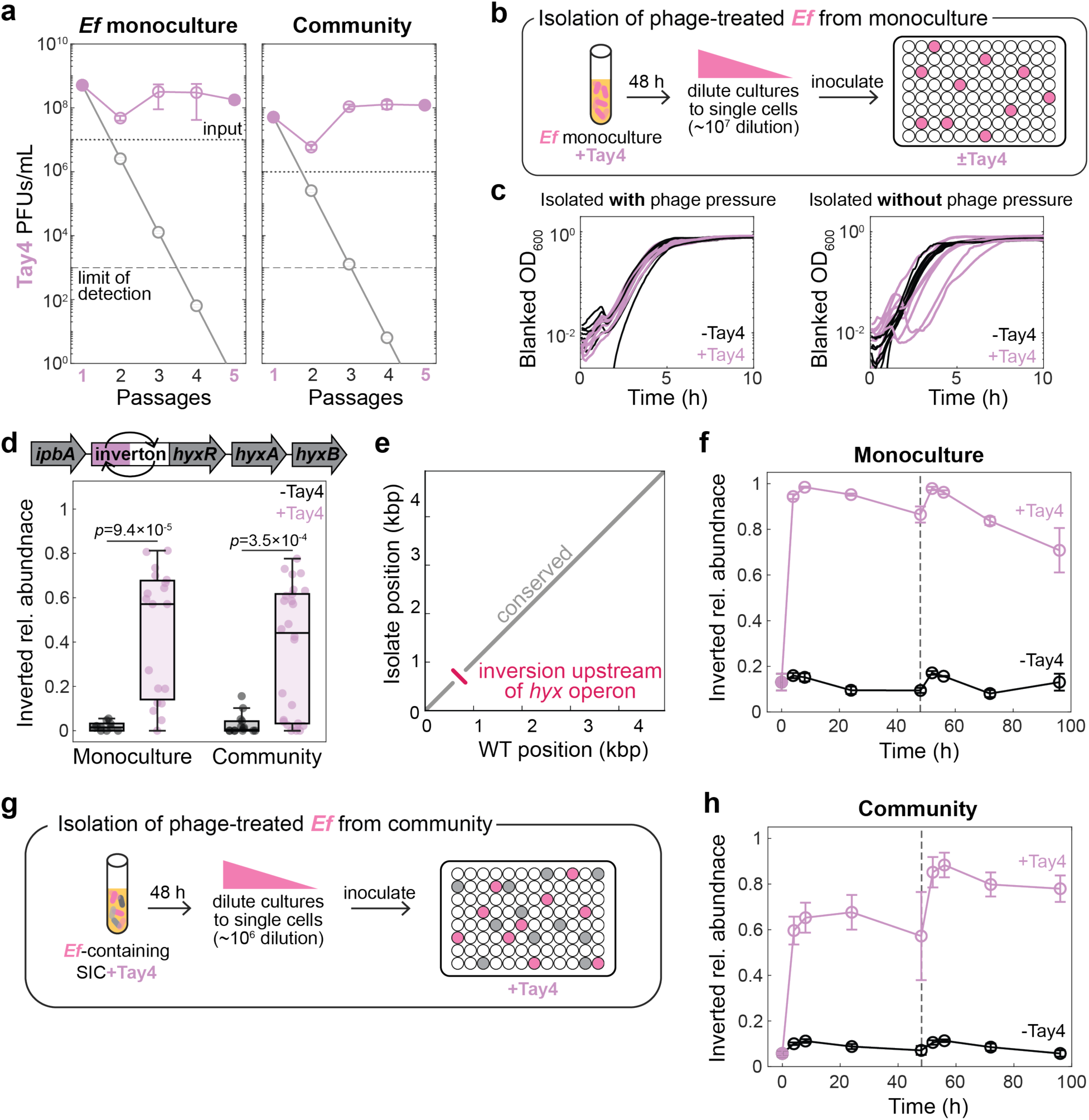
Phage resistance is mediated by reversible inversion upstream of the *hyx* operon and requires continual phage pressure. a) Tay4 titers during serial passaging of *Ef* in monoculture and in the MD2 community. Cultures were treated with Tay4 (MOI∼1) in passage 1, passaged without additional Tay4 in recovery passages 2–4, and treated again with Tay4 in passage 5. Phage titers were quantified after each passage by plaque assay (**Methods**). The gray line indicates the expected dilution trajectory (1:200 per passage) in the absence of phage propagation if the entire *Ef* population was resistant. Maintenance of high phage titers indicates continued replication on a susceptible *Ef* subpopulation. Points show mean±s.d. (*n*=3 biological replicates). Dotted line indicates input Tay4 concentration (∼10^7^ PFUs/mL) and dashed line indicates the limit of detection (∼10^3^ PFUs/mL). b) Schematic of single-cell isolation of phage-treated *Ef* from monoculture. Cultures were grown with Tay4 (MOI∼1), diluted to single-cell density (∼10^-7^), and distributed into 96-well plates with or without phage to assess stability of resistance. c) Growth dynamics of individual *Ef* isolates obtained under phage pressure (left, *n*=7) or without phage pressure (right, *n*=8). Isolates were passaged with (purple) or without (black) Tay4 (MOI∼1). Because the Tay4 titer was ∼10^8^ PFUs/mL at the end of the first treatment passage as in (**a**), it is highly unlikely that both a single cell and a phage particle were present in the same well. Resistance is associated with reduced lag time in the presence of phage, whereas isolates maintained without phage exhibited increased lag, consistent with reversion. d) Frequency of inversion of the promoter region upstream of the *hyx* operon. Short-read sequencing identified this inversion as the only structural variant consistently enriched during phage treatment in both monoculture and the MD2 community context (**Methods**). Box plots show distribution across biological replicates (*n*=10, 19, 14, and 27). Center line indicates the median; box limits, the interquartile range; whiskers, non-outlier range; points, individual replicates. *p*-values were calculated using two-sided Wilcoxon rank-sum tests. e) Synteny analysis of the local genomic region spanning the inverted promoter region of a representative phage-treated isolate mapped against the wild-type genome, showing inversion upstream of the *hyx* operon. Gray indicates conserved regions and red indicates the inverted segment. f) Dynamics of *hyx* promoter inversion during monoculture growth. *Ef* was grown with (purple) or without (black) Tay4 over two passages. Inversion frequency, measured by amplicon sequencing (**Methods**), increased rapidly upon phage exposure and decreased in its absence, indicating reversibility. Points show mean±s.d. (*n*=3 biological replicates). Dotted line marks passage transition (1:200 dilution). g) Schematic of single-cell isolation of phage-treated *Ef* from community growth. SICs were treated with Tay4 (MOI∼1), diluted to near single-cell density (∼10^-6^), and plated into 96-well plates with phage. *Ef*-containing wells were identified and sequenced to confirm identity and purity. h) Dynamics of *hyx* promoter inversion in *Ef* isolates derived from the MD2 community. As in monoculture (**f**), inversion frequency increased in the presence of Tay4 and decreased without phage, demonstrating that maintenance of resistance requires continual phage pressure. Points show mean±s.d. (*n*=3 biological replicates). Dotted line marks passage transition.

Consistent with this model, Tay4 failed to lyse or replicate on stationary-phase *Ef* cells (**Fig. S4c,d**), suggesting that selection for resistance occurs during exponential growth, whereas reversion to susceptibility occurs during stationary phase. Together, these results support a model in which phage resistance is dynamically maintained through reversible transitions, enabling coexistence of phage and host populations. These observations further suggest that resistance requires continual phage pressure, present during liquid growth but absent during colony-based isolation.

### Phage resistance is mediated by reversible inversion of the *hyx* operon

To identify the genetic basis of this reversible resistance, we developed a protocol to isolate *Ef* under continuous phage pressure (**Fig. 2b**, **Methods**). Cultures were diluted to near single-cell occupancy per well (**Fig. S4e**) and regrown in the presence or absence of Tay4, followed by re-exposure to phage. Isolates maintained under phage pressure retained resistance, whereas those grown without phage increased lag times upon re-exposure, consistent with partial reversion (**Fig. 2c**). This approach confirmed that phage pressure is required to maintain Tay4 resistance and enabled isolation of stable Tay4-resistant *Ef* populations.

We sequenced 19 resistant *Ef* isolates obtained under continuous phage pressure alongside 10 wild-type *Ef* isolates (**Fig. 2c, Methods**). No consistent point mutations or indels were detected across resistant genomes (**Fig. S5a**). Instead, the only structural variant enriched in resistant isolates was an inversion in the promoter region upstream of the *hyx* operon (**Fig. 2d,e**, **S5b**). This region lies downstream of *ipbA*, a gene encoding an invertase associated with phase variation in *E. coli*^49^. *ibpA* is upstream of the *hyx* operon, which contains genes involved in outer membrane structure, including O-antigen length^50^. Consistent with this, *hyxR* expression is controlled by bidirectional phase inversion of its promoter region, which impacts the survival of extraintestinal pathogenic *E. coli* in macrophages^51^. Time-resolved sequencing confirmed that this promoter undergoes dynamic inversion during Tay4 exposure, with partial reversion upon entry into stationary phase (**Fig.2f**, **Methods**). Together, these results indicate that resistance to Tay4 is mediated by reversible regulation of the *hyx* operon, with reversion in stationary phase generating a susceptible subpopulation that sustains phage propagation across passages.

We next asked whether the same mechanism operates in a community context. Using an adapted isolation protocol with continuous phage pressure (**Methods**), we obtained *Ef* isolates from Tay4-treated MD2 communities and sequenced 27 phage-treated and 14 susceptible isolates (**Fig. 2g**). As in monoculture, no consistent point mutations or indels were detected (**Fig. S5a**; **Methods**), and the only structural variant enriched in resistant isolates was the inversion upstream of the *hyx* operon (**Fig. 2d, S5b**). Time-resolved measurements confirmed enrichment of the inverted orientation during phage treatment in the community setting (**Fig. 2h**). These data suggest that resistance arises through the same reversible mechanism in both monoculture and community contexts, linking *hyx* regulation to the fitness trade-offs that reshape community composition during phage treatment.

### Successful species knockout using a phage cocktail and ecological constraints

While phage treatment can modulate community composition without complete species removal (**Fig. 1d**, **Fig. S3c**), defining the ecological role of *Ef* is much more straightforward to interrogate with its complete knockout. We therefore tested whether introducing a close competitor could suppress expansion of resistant *Ef* following Tay4 treatment. Metabolomics analysis revealed that *Kp* exhibited the highest niche overlap with *Ef* among isolated community members and the laboratory strain *Escherichia coli* MG1655 (**Fig. 3a**; **Methods**). However, addition of *Kp* during Tay4 treatment did not reduce the steady-state abundance of *Ef* across SICs (**Fig. 3b**, **S6a**), even at higher phage concentrations (**Fig. S6b**). Instead, *Kp* engrafted when niches were opened by *Ef* resistance (**Fig. S6a**), consistent with our observations above (**Fig. 1d-f**). These results indicate that ecological competition alone is insufficient to achieve complete *Ef* knockout and that knockout strategies should instead focus on phage efficacy.

**Figure 3:**
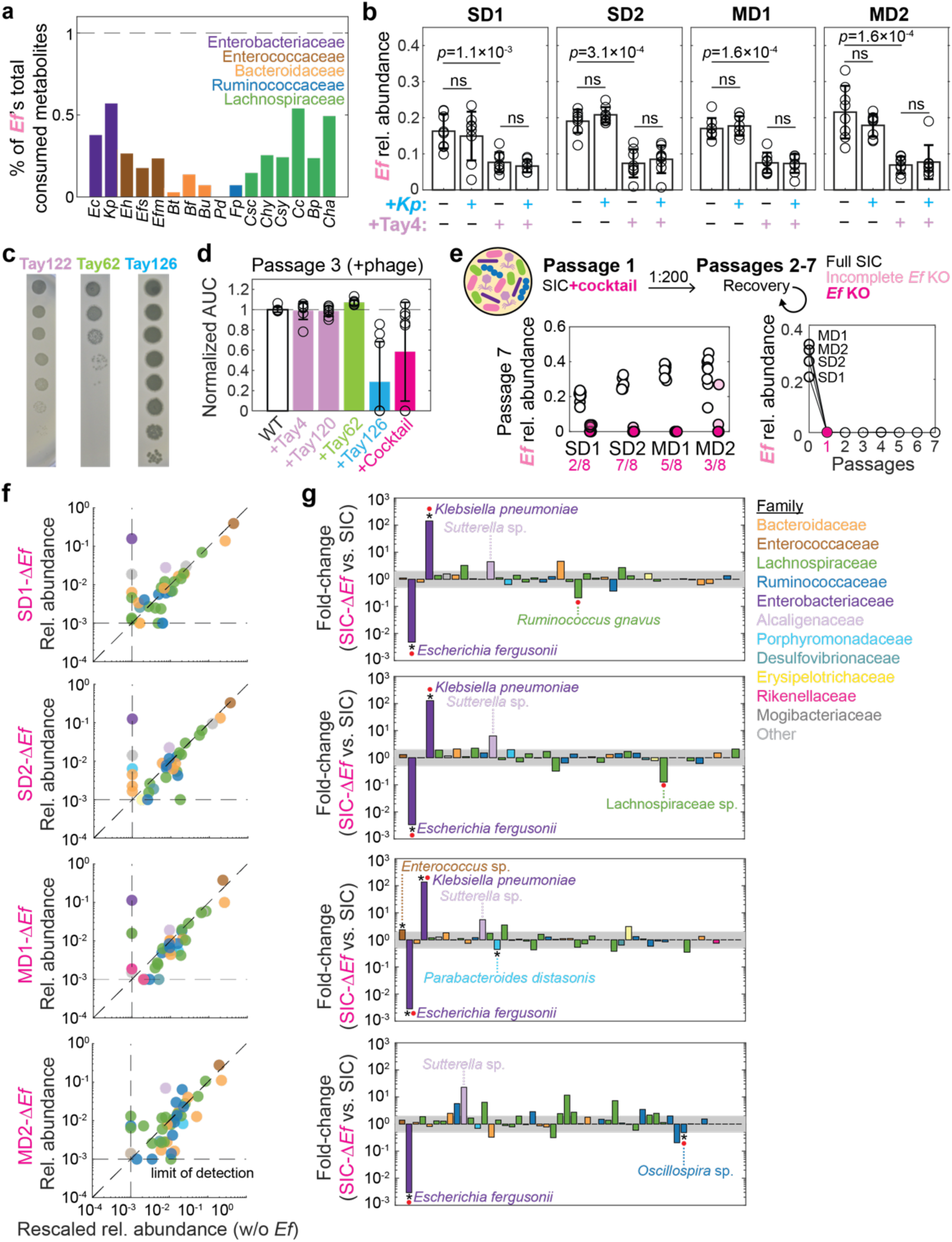
A phage cocktail enables *Ef* knockout with minimal disruption to community composition. a) Overlap in metabolite consumption between *Ef* and other community members. Bars show the fraction of metabolomic features consumed by *Ef* that are also depleted by each species during monoculture growth. Features were defined as consumed if they decreased >100-fold relative to fresh BHI, had peak area >5,000 in BHI, and were significantly depleted (two-sample Student’s *t*-test with Benjamini-Hochberg correction, adjusted *p*<0.05). *Kp* exhibited the greatest niche overlap with *Ef*. Bars are colored by family. Species abbreviations are in **Table S1**. b) Effect of competition and phage treatment on *Ef* abundance across SICs. Community were grown under four conditions: control, +*Kp*, +Tay4 (MOI∼1), and +*Kp* +Tay4. Competition with *Kp* did not enhance knockout efficacy. Bars show mean *Ef* relative abundance (*n*=8 biological replicates); error bars indicate ±s.d. *p*-values were calculated using two-sided Wilcoxon rank-sum tests (ns: not significant). c) Plaque assays of lytic phages Tay122, Tay62, and Tay126 on *Ef* lawns. d) Growth of *Ef* following treatment with individual phages or a phage cocktail. Normalized AUC (0–48 h) is shown for passage 3 growth following prior exposure and recovery (as in Fig. 1b). Points represent biological replicates (*n*=8); error bars indicate ±s.d. e) Phage cocktail enables *Ef* knockout across communities. Communities were treated with the phage cocktail in passage 1 and passaged without phage for recovery (passages 2–7). *Ef* relative abundance in passage 7 is shown for control communities (white) and phage-treated communities, stratified by whether *Ef* remained above (light pink) or below (magenta) the limit of detection. The fraction of successful knockouts (*n*=8 replicates per community) is indicated. f) Community composition following *Ef* removal after passage 7. Relative abundance of each ASV in Δ*Ef* communities is compared to the corresponding full community after rescaling to exclude *Ef*. Most ASVs lie near the line *y*=*x*, indicating minimal global perturbation. Points are colored by family; dashed lines indicate the limit of detection. g) Species-level changes following *Ef* removal. Bars show fold change in ASV abundance in Δ*Ef* communities relative to complete communities (mean across passages 5–7). The shaded region indicates 2-fold change. ASVs outside this region with significant changes (two-sample Student’s *t*-test with Benjamini-Hochberg correction, adjusted *p*<0.05) are marked with asterisks. ASVs that newly emerge above or drop below the limit of detection (10^-3^) are highlighted (red circles). Points are colored by family.

We next tested whether increasing phage diversity could overcome resistance^52–55^. We isolated three additional lytic phages from sewage sources and combined them with Tay4 at high titers to generate a four-phage cocktail (**Fig. 3c**; **Table S1; Methods**). All phages belong to the Caudoviricetes class and were genetically distinct (**Table S1**, **S3**). Similar to Tay4 alone, the cocktail did not inhibit the monoculture growth of 15 other strains isolated from these communities (**Fig. S2**), confirming its specificity. In monoculture, *Ef* developed resistance to Tay120 and Tay62 without detectable reduction in biomass yield (**Fig. 3d**), similar to Tay4. By contrast, the *Kagunavirus* phage Tay126 reduced biomass during resistance and achieved complete *Ef* knockout in some replicates, as did the full cocktail (**Fig. 3d**). Given our goal of robust community editing, we used the cocktail rather than a single phage to maximize knockout efficacy and reduce the likelihood that any single resistance mechanism would permit *Ef* recovery in community contexts.

Treatment of the four SICs with the phage cocktail (MOI∼100) resulted in successful knockout across all communities in at least some replicates (**Fig. 3e**). *Ef* decreased below the limit of detection and remained undetectable during subsequent passages (**Fig. 3e**), even after re-inoculation (**Fig. S7a**). Phage particles were also no longer detectable by plaque assays (**Fig. S7b**), confirming both species removal and phage clearance. Community biomass was unchanged following treatment (**Fig. S7c**), and composition was highly reproducible across replicates of a given community following *Ef* knockout (**Fig. S7d**). Hereafter, we denote representative replicate knockout communities as SD1-Δ*Ef*, SD2-Δ*Ef*, MD1-Δ*Ef*, and MD2-Δ*Ef*, and refer to untreated communities as complete communities.

Because *Ef* comprised ∼20%–40% of total community abundance prior to treatment, we expected that its removal might substantially alter community structure. However, community composition remained largely stable following *Ef* knockout, with most taxa unaffected after rescaling for the absence of *Ef* (**Fig. 3f,g**). The primary changes were confined to Proteobacteria: as observed during resistance (**Fig. 1d**). *Kp* expanded in communities where it was present, and a *Sutterella* species increased from low abundance to ∼5% across all SICs (**Fig. 3g**). Other changes were limited and community-specific but reproducible across passages (**Fig. S7e**), with no consistent expansion of non-Proteobacteria taxa. These results indicate that *Ef* occupies a largely orthogonal niche and that its removal primarily reshapes competition within a narrow taxonomic group, with modest effects on a small number of taxa including members of the *Ruminococcus*, *Enterococcus*, *Parabacteroides*, *Oscillospira*, and *Clostridium* genera **(Fig. 3g**, **Fig. S7e**).

We next asked whether *Ef* knockout could be improved by targeting communities in which *Ef* is at low abundance, as in stool. To do so, we re-derived communities from the same stool samples used to generate the SICs by adding the phage cocktail during initial inoculation into BHI (**Methods**). This approach resulted in *Ef* knockout in all replicates (**Fig. S8a**), a higher efficiency than treatment of established *Ef*-abundant SICs. Despite the absence of *Ef* during community assembly, overall diversity was unchanged (**Fig. S8b**), consistent with *Ef* occupying a largely orthogonal niche. Although compositional shifts were somewhat greater under these conditions (**Fig. S8c**), these results demonstrate that *Ef* can be robustly eliminated using phage cocktails and/or low-inoculum strategies.

### Reintroduction into ΔEf communities reveals privileged niche access of Ef

Because *Ef* can be removed from all four communities without substantially altering community composition, we used Δ*Ef* communities to test how *Ef* accesses its ecological niche. In a defined synthetic community, *Ef* reaches its steady-state abundance independent of its initial inoculum (**Fig. 1g**), consistent with preferential access to its niche^47^. We asked whether this property extends to complex communities. After re-establishing Δ*Ef* communities (**Methods**), we reintroduced *Ef* across a range of initial abundances and passages the communities to steady state (**Fig. 4a**). By the final passage, *Ef* abundance was independent of its starting inoculum, even when introduced at very low levels (<10 cells; **Fig. 4b**), and overall community composition was unchanged across inoculum conditions (**Fig. 4c**). These results indicate that *Ef* has privileged access to its niche in complex communities and that priority effects alone cannot explain the reduced in *Ef* fitness observed following phage treatment.

**Figure 4:**
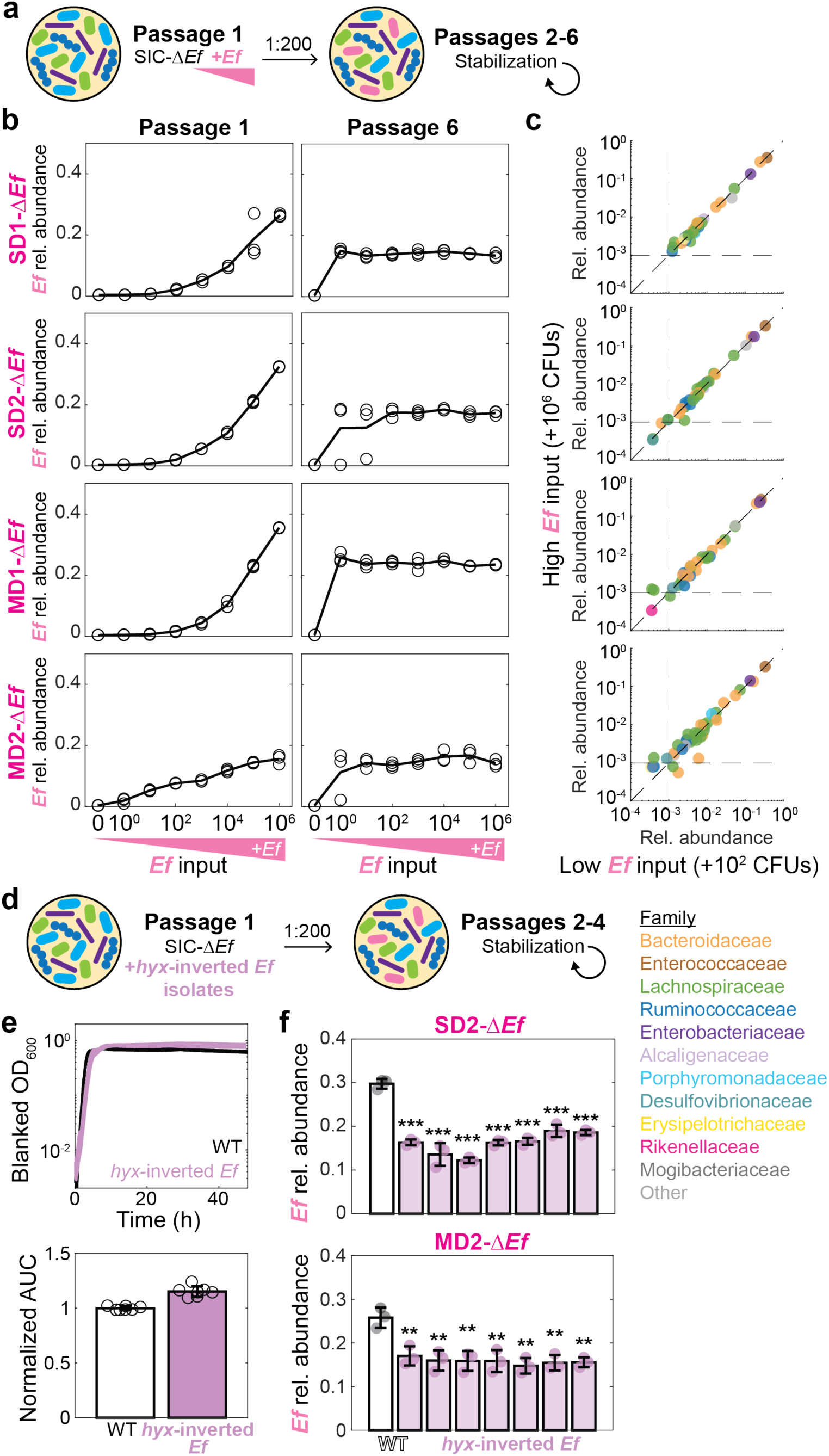
Knockout communities enable controlled reintroduction of *Ef* and confirm a community-specific fitness cost of phage resistance. a) Schematic of *Ef* titration into SIC-Δ*Ef* communities. *Ef* was added at the start of passage 1 across a range of initial abundances (10-fold dilutions), and communities were passaged (1:200 dilution) for five additional passages to reach steady state. b) *Ef* abundance converges to a steady-state value independent of initial inoculum. Relative abundance of *Ef* is shown after passage 1 (left) and passage 6 (right) for each SIC. Lines connect biological replicates (*n*=3), and points represent individual measurements. c) Community composition is insensitive to *Ef* initial abundance. Mean relative abundance of each ASV across biological replicates (*n*=3) at passage 6 is compared between communities initialized with high (∼10^6^ CFUs) or low (∼10^2^ CFUs) *Ef* input. Most ASVs lie near the line *y*=*x*, indicating minimal impact of initial *Ef* abundance on community structure. Points are colored by family; dashed lines indicate the limit of detection. d) Schematic of reintroduction of wild-type (WT) or phage-resistant (*hyx*-inverted) *Ef* isolates into SIC-Δ*Ef* communities. Strains were added at the start of passage 1 and cultures were passaged (1:200 dilution) for three additional passages to reach steady state. e) Growth curves (top) and normalized AUC (bottom) are similar between wild-type and *hyx*-inverted *Ef* isolates in monoculture, indicating no intrinsic fitness defect in isolation. Points represent biological replicates (*n*=7); bars indicate mean±s.d. f) Phage-resistant *Ef* isolates exhibit reduced fitness in community contexts. Relative abundance of wild-type and *hyx*-inverted *Ef* isolates following introduction into SD2-Δ*Ef* (top) and MD2-Δ*Ef* (bottom) communities. Data shown are after passage 4. Bars indicate mean±s.d. (*n*=3 biological replicates) and points represent individual replicates. Statistical significance was assessed relative to wild-type (two-sided Student’s *t*-test, **: *p*<0.01).

We next asked whether phage-resistant *Ef* exhibits a similar capacity to access its niche. We introduced seven Tay4-resistant isolates carrying the *hyx* promoter inversion (**Fig. 2c,d**) into the SD2-Δ*Ef* and MD2-Δ*Ef* communities under phage pressure (**Fig. 4d**). Although these isolates showed no fitness disadvantage relative to wild-type *Ef* in monoculture (**Fig. 4e**), they exhibited a marked reduction in abundance in community settings (**Fig. 4f**), comparable to Tay4-treated wild-type *Ef* (**Fig. 1d**). These results confirm that phage resistance incurs a fitness trade-off specifically in community contexts, impairing *Ef*’s access to its ecological niche. More broadly, these findings demonstrate that phage-mediated knockout enables controlled interrogation of species-level niche access by decoupling priority effects, inoculum size, and strain identity.

### Phage-mediated knockout reveals that *Ef* is necessary and sufficient for colonization resistance against *Salmonella* Typhimurium

The ability to remove a single species enables direct testing of its contribution to community function. In previous work, we found that SICs derived from stool of humanized mice before ciprofloxacin treatment were resistant to invasion by Enterobacteriaceae member *Salmonella* Typhimurium (*S*Tm), whereas SICs derived from the same mice after ciprofloxacin treatment were highly susceptible^38^, consistent with *in vivo* observations that antibiotic treatment can increase vulnerability to pathogen invasion^56,57^. Because ciprofloxacin eliminated multiple taxa (including *Ef*)^38^, it remained unclear whether resistance to *S*Tm reflected removal of *Ef* specifically or disruption of a broader community state. Given that phage-mediated knockout of *Ef* minimally perturbs overall community composition (**Fig. 3f,g**), we used Δ*Ef* communities to test whether *Ef* is required for resistance to *S*Tm.

We challenged complete and Δ*Ef* communities with *S*Tm across a range of inoculum ratios and quantified *S*Tm abundance after passaging to steady state (**Fig. 5a**; **Methods**). *S*Tm densities were ∼100-fold higher in Δ*Ef* communities than in complete communities (**Fig. S9a**), and 16S sequencing showed that *S*Tm reached ∼10%-15% relative abundance in Δ*Ef* communities, independent of initial *S*Tm inoculum size, but remained below the limit of detection when *Ef* was present (**Fig. 5b**). *S*Tm coexisted with other Proteobacteria such as *Kp* in Δ*Ef* communities (**Fig. 5b**), suggesting that *Ef* competitively excludes *S*Tm within a shared niche. The abundance of most other taxa was largely unchanged (**Fig. S9b**). These results demonstrate that *Ef* is necessary for preventing *S*Tm expansion.

**Figure 5:**
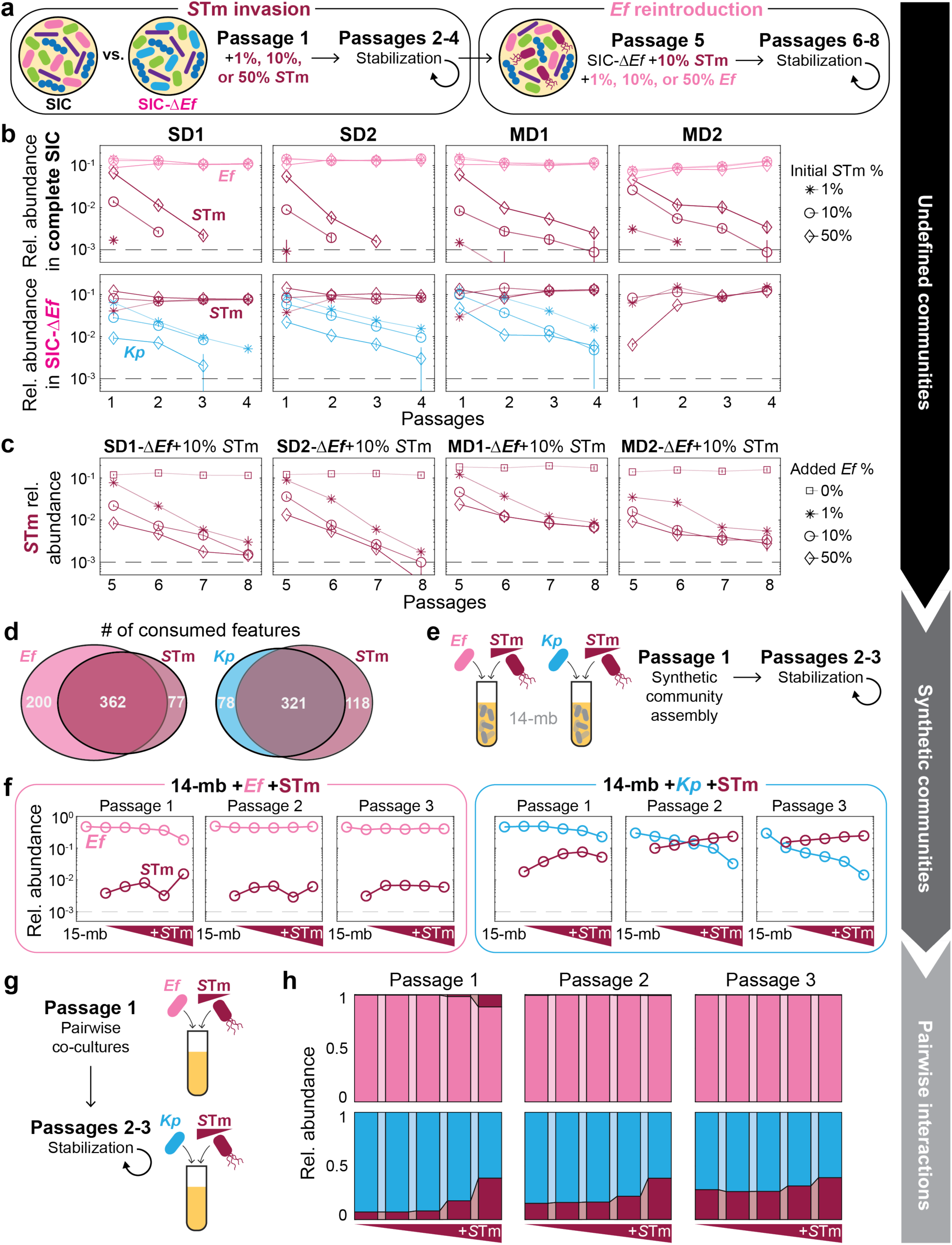
*Ef* is necessary and sufficient for community resistance to *Salmonella* invasion. a) Schematic of *S*Tm challenge. *S*Tm was introduced into complete or Δ*Ef* communities at 1%, 10%, or 50% initial abundance and passaged for stabilization four times. *Ef* was then introduced at defined abundances (0%, 1%, 10%, or 50%) into Δ*Ef* communities challenged with 10% *S*Tm, and communities were further passaged to assess displacement of established *S*Tm (passages 5-8). b) Dynamics of Enterobacteriaceae ASVs during *S*Tm invasion (passages 1–4) into complete (top) or SIC-Δ*Ef* (bottom) communities. Relative abundance of *Ef* (pink), *S*Tm (maroon), and *Kp* (blue) are shown for different *S*Tm inocula (1%, 10%, 50%). *Kp* abundance was below the limit of detection in the complete communities. Points represent mean±s.d. (*n*=3 biological replicates). Dashed line indicates the limit of detection. c) *Ef* reintroduction displaces established *S*Tm. *S*Tm abundance is shown during passages 5–8 following reintroduction of *Ef* into SIC-Δ*Ef* communities initially challenged with 10% *S*Tm. *Ef* was added at 0%, 1%, 10%, or 50%. Points represent mean±s.d. (*n*=3 biological replicates). Dashed line indicates the limit of detection. d) Overlap in metabolite consumption between *Ef*, *S*Tm, and *Kp*. Venn diagrams show the number of metabolomic features uniquely or jointly depleted during monoculture growth. Features were defined as consumed if they decreased >100-fold relative to fresh BHI, had peak area >5,000 in BHI, and were significantly depleted (two-sample Student’s *t*-test with Benjamini-Hochberg correction, adjusted *p*<0.05). e) Schematic of synthetic community invasion assays. Communities were assembled from isolates (equal OD mixture) with *S*Tm introduced across a range of inputs (10^5^-10^9^ CFUs/mL) into 15-member communities containing either *Ef* or *Kp*, followed by stabilization over two additional passages. f) *S*Tm invasion dynamics in synthetic communities. Left: *Ef*-containing communities; right: *Kp*-containing communities. Relative abundances of *Ef* (pink), *Kp* (blue), and *S*Tm (maroon) are shown across three passages. In the presence of *Ef*, *S*Tm was excluded, whereas *S*Tm persisted in *Kp*-containing communities. Representative experiment shown. g) Schematic of pairwise competition assays between *S*Tm and *Ef* or *Kp* across a range of initial *S*Tm abundances. *Ef* or *Kp* was added at 10^9^ CFUs/mL, and *S*Tm was added at 10-fold dilutions ranging from 10^5^-10^9^ CFUs/mL at the beginning of passage 1. Co-cultures were allowed to stabilize for two additional passages. h) Pairwise competition dynamics. Relative abundance of *S*Tm (maroon) with *Ef* (pink) or *Kp* (blue) are shown across three passages*. Ef* outcompeted *S*Tm, whereas *Kp* coexisted with *S*Tm. Representative experiment shown.

To test whether *Ef* is sufficient to restore colonization resistance, we introduced *Ef* into *S*Tm-invaded Δ*Ef* communities after initial stabilization (**Fig. 5a**). Across all inoculation ratios, addition of *Ef* reduced *S*Tm abundance by ∼100-fold in both absolute (**Fig. S9c**) and relative abundances (**Fig. 5c**). These results indicate that *Ef* is sufficient to suppress *S*Tm engraftment.

We next investigated the competitive basis of this interaction. Metabolomics profiling revealed that *S*Tm shares a larger fraction of its niche with *Ef* (∼82%) than with *Kp* (∼73%) (**Fig. 5d**). Consistent with this, *S*Tm was robustly outcompeted in a synthetic community containing *Ef* across all inoculation densities but coexisted when *Ef* was replaced with *Kp* (**Fig. 5e,f**). In this setting, *S*Tm abundance scaled with its initial inoculum when co-cultured with *Kp*, consistent with approximately neutral competition^58^ and their coexistence in Δ*Ef* communities (**Fig. 5b**). Pairwise co-culture experiments further showed that *S*Tm coexists with *Kp* but is driven to near the limit of detection by *Ef*, even when initially introduced at higher abundance (**Fig. 5g,h**). These results indicate that *Ef*-mediated colonization resistance arises from strong competitive exclusion of *S*Tm within a shared nutritional niche, and that knockout communities can be used to elucidate the role of a single species and its ecological interactions within undefined communities.

### Replacement with genome-wide mutant library reveals that outer membrane integrity determines *Ef* fitness in communities

To identify the genetic determinants of *Ef* fitness in community contexts, we leveraged phage-mediated knockout to replace *Ef* with a genome-wide library of mutants. We constructed a pooled library of barcoded transposon insertions comprising 1,025,611 unique mutants, providing near-saturating coverage of the *Ef* genome. The library was introduced into each of the four Δ*Ef* communities and grown in parallel in monoculture (**Fig. 6a**), and changes in mutant abundance were quantified by barcode sequencing^59^ (BarSeq; **Methods**; **Table S4**).

**Figure 6:**
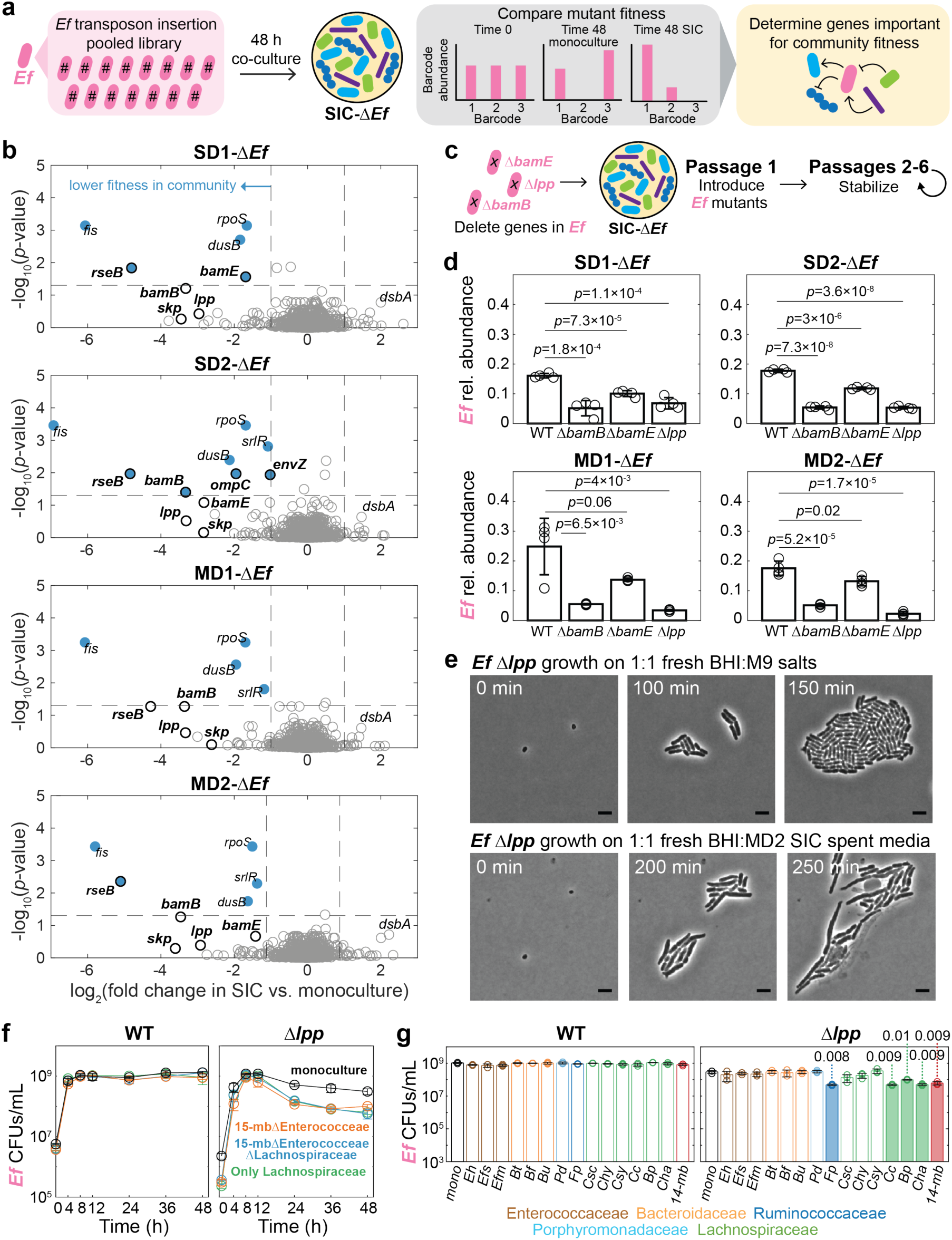
Replacement of *Ef* with a genome-wide mutant library reveals the importance of outer membrane integrity for *Ef* fitness in a community. a) Schematic of pooled transposon mutant fitness profiling. A barcoded *Ef* transposon insertion library was grown in monoculture or co-cultured with *Ef*-knockout communities (SIC-Δ*Ef*). Mutant fitness was quantified by changes in barcode abundance after 48 h to identify genes specifically required for growth in communities. b) Genome-wide identification of genes required for community fitness. Mutant fitness is shown as log_2_ fold change in abundance in SICs relative to monoculture. Each point represents a gene, aggregated across insertion mutants. Genes with significantly reduced fitness in communities (two-sample Student’s *t*-test with Benjamini-Hochberg correction, adjusted *p*<0.05, *n*=3 biological replicates) are highlighted in blue. Vertical dashed lines indicate 2-fold increase/decrease. Horizontal dashed line indicates significance threshold. Genes involved in outer membrane assembly and structure are labeled with bold text and a black outline. c) Schematic for validation of candidate genes. Wild-type *Ef* and deletion mutants were individually introduced into SIC-*ΔEf* communities and passaged to steady state. d) Reduced fitness of outer membrane in communities. Relative abundance of wild-type *Ef* and deletion mutants (Δ*bamB*, Δ*bamE*, Δ*lpp*) after passage 6 in SIC-Δ*Ef* communities. Bars show mean±s.d. (*n*=4 biological replicates), with individual replicates overlaid. *p*-values were calculated using two-sample Student’s *t*-tests. e) Time-lapse microscopy of Δ*lpp* cells. Top: growth on 50% BHI-50% M9 salts showing normal morphologies (**Movie S2**). Bottom: growth in 50% BHI-50% MD2-Δ*Ef* spent medium showing elongation, swelling, and lysis (**Movie S4**). Scale bars: 5 µm. f) Growth dynamics of wild-type (WT) and Δ*lpp Ef* in monoculture and community contexts in BHI-S. CFU measurements show that Δ*lpp* reaches similar maximum density as wild type but undergoes rapid decline during stationary phase when co-cultured with community members. *Ef* was grown in a 15-mb community without Enterococcaceae species (–*Eh*, –*Efs*, –*Efm*), a 15-mb community without Enterococcaceae or Lachnospiraceae species (–*Eh*, –*Efs*, –*Efm,* –*Csc*, –*Chy*, –*Csy*, –*Cc*, –*Bp*, –*Cha*), and a synthetic community consisting only of Lachnospiraceae species (*Csc*, *Chy*, *Csy*, *Cc*, *Bp*, *Cha*). Points show mean±s.d. (*n*=3 biological replicates). g) Pairwise interactions reveal multiple sources of antagonism. CFU measurements of wild-type and Δ*lpp Ef* after 48 h co-culture with individual community members. Bars show mean±s.d. (*n*=3 biological replicates), with individual replicates overlaid. Significant differences relative to monoculture (two-sample Student’s *t*-test with Benjamini Hochberg correction, adjusted *p*<0.05) are indicates. Colors denote taxonomic family.

Comparing mutant fitness in communities to monoculture revealed a small set of genes whose disruption specifically reduced fitness in community contexts (**Fig. 6b**; **Methods**). These genes were largely shared across all communities and exhibited consistent fitness defects, suggesting a general mechanism underlying community-specific fitness. Mutants in the global regulator *fis*^60^ and stress response genes such as *rpoS*^61^ showed strong fitness decreases (**Fig. 6b**), consistent with broad roles in cellular physiology. Strikingly, genes involved in outer membrane protein biogenesis (*bamB*, *bamE, skp*, *ompC, envZ*), outer membrane–cell wall connections (*lpp*), or envelope stress responses (*rseB*) were among the most consistently depleted in communities relative to monoculture (**Fig. 6b**). In contrast, mutations in genes flanking the *hyx* operon did not exhibit decreased fitness in community settings, consistent with resistance arising from phase-variable activation of the operon rather than gene disruption^49^. Together, these results identify outer membrane integrity as a key determinant of *Ef* fitness in microbial communities.

### Outer membrane defects sensitize *Ef* to community-derived antagonism

To determine how outer membrane defects reduce fitness in communities, we constructed deletion mutants in representative genes identified in the screen (*bamB*, *bamE*, and *lpp*; **Methods**). These mutants exhibited increased outer membrane permeability, as evidenced by sensitivity to SDS/EDTA (**Fig. S10a,b**) and, for Δ*bamB*, to vancomycin (**Fig. S10c**). When introduced into Δ*Ef* communities (**Fig. 6c**), all mutants exhibited reduced abundance relative to wild-type *Ef* after a single passage (**Fig. S10d**), and fitness defects persisted at steady state (**Fig. 6d**). In monoculture, the mutants reached similar maximum densities as wild-type *Ef*, but Δ*bamB* and Δ*lpp* exhibited reduced final yield (**Fig. S10e-g**), suggesting increased sensitivity to late–stationary-phase stress. Co-culture with both the MD2-Δ*Ef* SIC and the 14-member synthetic community further reduced mutant fitness relative to wild-type *Ef* (**Fig. S10h,i**), indicating that defects are robust across community contexts and likely arises from interactions with other community members.

Given the established role of the outer membrane in protecting against environmental stresses^62^, we hypothesized that defects in envelope integrity sensitize *Ef* to toxic molecules produced by other community members. To test this, we used anaerobic time-lapse microscopy (**Methods**) to examine wild-type *Ef* and Δ*lpp* cells growth in the presence of MD2-Δ*Ef* community spent medium. While wild-type *Ef* growth was unaffected, Δ*lpp* cells exhibited rapid swelling, loss of shape, and lysis within hours (**Fig. 6e**, **S11a,b**; **Movies S2–S5**), indicating heightened sensitivity to community-derived factors. This effect was abolished by heat treatment of the spent medium (**Fig. S11c**; **Movie S6**; **Methods**), suggesting that the responsible factors are heat-labile. Together, these results demonstrate that compromised outer membranes sensitize *Ef* to toxic molecules produced by the community.

To further characterize this interaction, we examined how growth conditions influence the sensitivity of outer membrane mutants. All Δ*Ef* SICs had a final pH of ∼7.0, arguing against environmental pH as the cause of the Δ*lpp* morphological defects. When grown in PBS-diluted spent medium, Δ*lpp* cells exhibited reduced growth without the pronounced morphological defects observed under growth-permissive conditions (**Fig. S11d**; **Movie S7**), suggesting that antagonistic effects are amplified during rapid growth. Together, these results indicate that the fitness defect of outer membrane mutants depends on both exposure to community-derived factors and the physiological state of the cells.

Because the Δ*lpp* mutant exhibited reduced fitness in both undefined SICs and the defined 14-member synthetic community (**Fig. S10h,i**), we leveraged the isolate-resolved community to identify species that antagonize Δ*lpp* growth. To distinguish whether the fitness defect arises from nutrient competition or direct antagonism, we compared growth dynamics of wild-type and Δ*lpp Ef* in monoculture and in community co-culture. Both strains reached similar maximum abundance during exponential growth, suggesting that nutrient competition is not the primary driver of the fitness defect, but Δ*lpp* cells exhibited a pronounced decline during stationary phase (**Fig. 6f**), indicating increased cell death. This defect was exacerbated in the presence of the community (**Fig. 6f**), consistent with enhanced sensitivity to community-derived factors. The timing of this decline coincided with saturation of the Lachnospiraceae and Bacteroidaceae families^41^, suggesting that antagonistic interactions emerge during late growth phases. Together, these results indicate that the reduced fitness of Δ*lpp* is not explained by impaired nutrient acquisition but instead reflects increased susceptibility to community-induced cell death.

Genomic analysis revealed that 12 of the 14 species encode ribosomally synthesized and post-translationally modified peptides (RiPPs; **Fig. S11e**; **Methods**), which can target bacterial cell envelopes^63^. Pairwise co-culture experiments showed that several species, including Lachnospiraceae members *Clostridium clostridioforme*, *Clostridium hathewayi*, and *Blautia producta* and the Oscillospiraceae *Flavonifractor plautii*, recapitulated the fitness defect of Δ*lpp* observed in complete communities (**Fig. 6g**). All of these species have the capacity to synthesize RiPPs (**Fig. S11e**), linking the observed antagonism to potential envelope-targeting mechanisms. These results indicate that multiple community members produce factors that selectively inhibit *Ef* with compromised outer membranes.

Our pairwise co-culture results indicate that multiple species secrete metabolites that antagonize *Ef Δlpp*, potentially through more than one mechanism. However, these species did not fully recapitulate the effects observed in the full MD2 community. Consistent with this, *Ef* Δ*lpp* cells did not exhibit the pronounced morphological defects seen with MD2 spent medium when grown with spent medium from the 14-member synthetic community (**Fig. S11f**; **Movie S8**). Together, these findings suggest that additional community members and/or factors contribute to the full antagonistic phenotype, which can manifest as cell death in stationary phase and morphological defects. These data demonstrate that phage-mediated knockout of a single species can reveal complex, multi-species interactions that shape bacterial fitness in microbial communities.

## Discussion

In this study, we demonstrate that phage-mediated knockout of a single bacterial species from complex gut microbial communities can reveal otherwise obscured ecological interactions, reshape community composition, and modulate community-level functions. By targeting *Ef*, a dominant member with few apparent competitors, we found that even incomplete phage-mediated depletion could restructure communities through fitness trade-offs with taxa emerging from very low abundance (**Fig. 1d**). Complete knockout was achieved using a phage cocktail and by targeting populations at low abundance to overcome resistance (**Fig. 3e, S8a**). Removal of *Ef* altered the competitive landscape, enabling expansion of specific Proteobacteria species and revealing its essential role in colonization resistance against *Salmonella* (**Fig. 5a-c**). In contrast, non-Proteobacteria species were largely unaffected, consistent with limited niche overlap (**Fig. 3g**) and potential functional redundancy between *Kp* and *Ef* (e.g., heme production^47^). The preservation of overall community structure highlights the utility of phage-mediated knockout as a tool for mechanistic interrogation of microbial ecosystems. At the genetic level, we identified outer membrane integrity as a key determinant of *Ef* fitness during phage exposure (**Fig. 2d**, **4f**) and in community contexts (**Fig. 6b-d**), and we uncovered widespread interspecies antagonism (**Fig. 6e-g**) that is masked in standard co-culture assays. Together, these findings establish phages not only as therapeutic agents but also as ecological tools for dissecting microbial assembly, competition, and cooperation.

Our results further show that the ecological landscape of a community critically shapes the efficacy of phage-based editing strategies. *Ef* persists despite targeted phage treatment due to its privileged access to a metabolic niche: even a single resistant cell, once established, can repopulate the community. This resilience highlights a broader challenge for phage-mediated manipulation, as species with strongly partitioned resource use^47^ may be intrinsically difficult to eliminate regardless of niche size. Consistent with this, introducing a close competitor such as *Kp* was insufficient to displace resistant *Ef* (**Fig. 3b**, **S6a-b**), although such strategies may be more effective for species with less exclusive niches. Together, these findings emphasize the importance of understanding competitive structure and improving phage efficacy, through cocktails, engineered host range, or timing interventions to coincide with population bottlenecks.

Our study also reveals the dynamic nature of phage resistance in community settings. Given the large population sizes typical of gut communities, resistance is likely inevitable for abundant species due to inherent genetic heterogeneity. However, targeting *Ef* at low abundance substantially improved knockout efficiency (**Fig. S8a**), suggesting that phage-based strategies may be most effective when combined with perturbations (e.g., antibiotics) that transiently reduce target populations. We show that *Ef* resistance depends on continual phage pressure (**Fig. 2c**) and is mediated by reversible promoter inversion (**Fig. 2d-h**), enabling reversion to susceptibility in the absence of selection. This mechanism complements prior work demonstrating that invertible elements enable rapid and reversible adaptation in gut bacteria^64,65^. The reversibility of resistance has important implications for isolating resistant strains (**Fig. 2b,g**), and understanding resistance mechanisms is critical for creating opportunities for iterative selection of phages that target resistant subpopulations. More broadly, these findings motivate studies of species-specific phage responses across diverse microbial communities. Do all species exhibit similar resistance dynamics and niche protection, or are some more vulnerable to competition? Are resistance mechanisms typically associated with metabolic trade-offs, and are they reversible in the absence of phage pressure? Addressing these questions will help define the constraints governing phage-mediated community editing and establish its generalizability as a strategy for precision microbiome modulation.

Beyond ecological insight, our results suggest practical strategies for microbiome engineering and therapeutic intervention. Targeted phage treatment could be used to deplete specific taxa from complex communities, such as fecal microbiota transplant material, to reduce the risk of transmitting undesirable microbes. However, removal of a dominant species can also enable expansion of previously undetectable taxa, with potentially unpredictable consequences for community function, highlighting the importance of understanding the ecological impacts of species knockout. Phage-based editing may also facilitate the isolation of low-abundance organisms by suppressing dominant competitors, thereby improving recovery of rare but functionally important species. Together, these findings highlight both the potential and the challenges of using phages to precisely manipulate microbial communities.

More broadly, our work extends the logic of genetic perturbation to microbial ecology. Just as gene knockouts transformed our understanding of cellular systems, targeted species removal provides a framework for dissecting ecological interactions in complex communities. By selectively removing a single species and measuring the resulting shifts in composition and function, it becomes possible to assign ecological roles and dependencies. While nutrient competition is a central axis of microbial interaction, our findings highlight the importance of additional forces, including antagonistic interactions and envelope-targeting toxins, that shape community structure. The sensitivity of *Ef* outer membrane mutants to community-derived factors underscores the extent of interspecies antagonism within gut ecosystems and suggests that bacterial fitness is shaped by overlapping, multi-species pressures. Although our experiments were conducted *in vitro* under controlled conditions, they establish a foundation for future studies in more complex host-associated environments. Extending this approach to diverse communities, species, and engineered phages will be essential to fully realize phage-based editing as both a discovery platform and a strategy for precision microbiome modulation.

## Methods

### Bacterial strains and culturing conditions

All culturing was performed in an anaerobic chamber (Coy Labs) unless otherwise stated. Media and materials were equilibrated under anaerobic conditions for at least 48 h prior to use. Standard passaging consisted of a 1:200 dilution of saturated cultures into fresh medium followed by 48 h of growth at 37 °C.

Isolates, including *Ef*, members of the 15-member synthetic community, and *Kp,* were obtained by plating SICs from humanized mouse fecal samples on agar plates containing various complex media, as previously described^38,41^. *Escherichia coli* MG1655 was used for metabolomics comparison, and *Salmonella enterica* serovar Typhimurium SL1344 was used for invasion experiments.

Isolates were stored as 25% glycerol stocks at –70 °C. Frozen stocks were streaked onto BHI-blood agar plates (5% defibrinated horse blood in 1.5% (w/v) agar) and colonies were inoculated into 3 mL of Brain Heart Infusion (BHI, BD #2237500) or BHI supplemented with hemin and vitamin K1 (BHI-S) for isolate and synthetic community experiments. Cultures were grown anaerobically for 48 h at 37 °C.

For monoculture experiments, 1 μL of saturated culture was inoculated into 200 μL of fresh medium and grown anaerobically for 48 h at 37 °C. For synthetic community experiments, isolates were normalized by optical density at 600 nm (OD_600_), mixed at equal abundance, and inoculated (1 μL of the mixture into 200 μL of medium) prior to incubation anaerobically for 48 h at 37 °C. SICs were revived from 25% glycerol stocks by inoculating 5 μL into 1 mL of BHI and incubating anaerobically for 48 h at 37 °C. To minimize physiological variation due to from freeze-thaw cycles, SICs were passaged three times (1:200 dilution every 48 h) prior to experiments.

Phage-treatment experiments were conducted in BHI supplemented with 1 mM CaCl_2_. Phage lysate (1 μL) was added to cultures to achieve the desired MOI, calculated based on initial phage titer and estimated cell density. The number of *Ef* CFUs/mL in the communities was ∼10^8^, compared to ∼10^9^ in monoculture (**Fig. S3a**), and phage titer was adjusted accordingly to maintain equal MOI across conditions. Cultures were passaged every 48 h in 200 μL in clear, flat-bottom 96-well plates (Greiner Bio-One #655161) sealed with breathable membranes (Excel Scientific #BS-25) and lids to limit evaporation. Following growth, cultures were covered with a foil seal (Thermo Scientific #232698) and stored at –70 °C for downstream analyses.

### Growth measurements and analysis

Biomass over time was quantified by OD_600_ using an Epoch 2 plate reader (Biotek) operated within an anaerobic chamber. Measurements were performed in clear, flat-bottom 96-well plates. Background OD_600_ was measured for each well prior to inoculation and subtracted from subsequent readings. Cultures were inoculated by adding 1 μL of stationary-phase culture to 199 µL of BHI or BHI+CaCl_2_ immediately prior to measurement. Plates were sealed with transparent membranes (Excel Scientific #STR-SEAL-PLT) perforated above each well (∼0.5 mm diameter) to allow gas exchange. OD_600_ was measured every ∼5 min at 37 °C with continuous shaking. Growth was quantified as the area under the curve (AUC) by numerical integration of OD_600_ measurements from 0–48 h using the trapezoidal rule (Matlab *trapz* function).

### Phage isolation and purification

Bacteriophages were isolated from environmental samples collected from multiple wastewater treatment facilities (**Table S1**) using previously described methods^24^. Samples were centrifuged at 4,000*g* for 10 min at room temperature to remove solids, and supernatants were sequentially filtered through 0.45 μm and 0.22 μm polyethersulfone (PES) filters (Thermo Scientific #725-2445, #725-2520). Filtrates were concentrated up to 500-fold using 100-kDa molecular weight cutoff PVDF centrifugal filters (Millipore Sigma #UFC910024).

Phages were screened using a soft agar overlay assay. Briefly, 10-100 μL of concentrated filtrate were mixed with 0.5 mL of an overnight culture of *Ef* and 4.5 mL of molten BHI top agar (0.35%), then poured onto BHI agar plates (1.5% agar). Plates were incubated aerobically overnight at 37 °C. Individual plaques were picked and resuspended in 50 μL of phage buffer (10 mM Tris-HCl [pH 7.5], 10 mM MgSO_4_, 68 mM NaCl). Plaque purification was performed through at least three rounds to obtain isolated plaques.

High-titer phage stocks were generated by flooding plates exhibiting near-confluent lysis with 5 mL of phage buffer and incubating for at least 2 h at room temperature. The lysate was clarified by centrifugation (4,000*g*, 10 min) and filter-sterilized through a 0.22 μm PES filter. Phage stocks were stored at 4 °C.

### High-titer phage stock preparation

For phage-based community editing, concentrated phage stocks were prepared to enable high MOI. A single colony of *Ef* was inoculated into 5 mL of BHI and grown overnight with shaking at 37 °C. Overnight cultures were diluted 1:10 into 100 mL of fresh BHI supplemented with 1 mM CaCl_2_ and grown with shaking at 37 °C for ∼1 h to reach exponential phase. Phage was then added at MOI∼1 and cultures were incubated overnight at 37 °C.

Cultures were clarified by centrifugation at 4,000*g* for 10 min, and the supernatant was filter-sterilized through a 0.22 μm PES filter. The filtrate was concentrated up to 100-fold using 100-kDa PVDF molecular weight cutoff filters. Phage titers were determined by plaque assay using 10-fold serial dilutions on soft agar overlays of *Ef*. Concentrated phage stocks were stored at 4 °C.

### Phage DNA extraction

Phage DNA was extracted from high-titer lysates using a phenyl-chloroform extraction protocol. To remove contaminating bacterial DNA, 175 μL of phage lysate were treated with 5 μL of DNase I and 20 μL of reaction buffer (Qiagen #79254) at room temperature for 1 h. Capsid proteins were digested by adding 12.5 μL of 1 M Tris-HCl, 12.5 μL of 0.5 M EDTA, 12.5 μL of 20% SDS, and 5 μL of 20 mg/mL proteinase K (Thermo Scientific #EO0492), in that order, followed by incubation at 50 °C for 1 h.

DNA (200 µL) was purified by extraction with an equal volume of phenol/chloroform/isoamyl alcohol (Invitrogen #15593-049) using phase-lock gel tubes (Quantabio #2302830), followed by centrifugation at maximum speed for 5 min (Eppendorf Centrifuge #5424). The aqueous phase was re-extracted with elution buffer (10 mM Tris-HCl [pH 8.5]) and centrifuged to maximize DNA recovery. The recovered aqueous phase was then extracted with an equal volume of chloroform/isoamyl alcohol (Thermo Scientific #327155000) to remove residual phenol.

The aqueous phase (∼380 µL) was carefully transferred to a fresh tube for ethanol precipitation. DNA was precipitated by addition of ammonium acetate (final concentration 0.75 M) and 2.5X volumes of ice-cold 100% ethanol, followed by centrifugation at 4 °C (Eppendorf Centrifuge #5417R) for 20 min. Pellets were washed twice with 70% ethanol, air-dried, and resuspended in 50 μL of elution buffer. DNA was stored at 4 °C.

### Phage treatment and cocktail preparation

All phage-treated cultures were grown in BHI supplemented with 1 mM CaCl_2_. Phage lysates were diluted in phage buffer immediately prior to use to achieve the desired MOI. For heat inactivation, phage lysates were incubated at 95 °C for 10 min.

Phage cocktails were prepared by combining equal volumes of high-titer phage stocks immediately prior to use. Unless otherwise indicated, the cocktail consisted of Tay4 (∼10^9^ PFUs/mL), Tay62 (∼10^9^ PFUs/mL), Tay122 (∼10^9^ PFUs/mL), and Tay126 (∼10^10^ PFUs/mL).

### Colony forming unit (CFU) quantification

CFUs were quantified by 10-fold serial dilution of cultures in sterile 1X PBS (Thermo Scientific #AM9624), followed by spotting of 3 μL of each dilution onto agar plates. Plates were incubated at 37 °C and colonies were counted after incubation. For *Ef* monocultures, dilutions were plated on BHI agar (1.5%) and incubated aerobically overnight. For total community CFUs, dilutions were plated anaerobically on BHI agar supplemented with 5% defibrinated horse blood and incubated anaerobically at 37 °C for 48 h. To quantify *Ef* CFUs from communities, dilutions were plated aerobically on BHI agar and incubated overnight aerobically at 37 °C. *Ef* colonies were distinguished from other facultative anaerobes based on colony morphology, including larger colony size.

### Plaque forming unit (PFU) quantification

Phage titers were quantified by plaque assay. Phage stocks or filter-sterilized phage-treated cultures were serially diluted 10-fold in sterile phage buffer. Agar plates were prepared by mixing 0.5 mL of an overnight culture of *Ef* with 4.5 mL of molten BHI top agar (0.35%) and overlaying this mixture onto BHI agar (1.5%) plates. After solidification, 3 μL of each dilution were spotted onto the surface. Plates were incubated aerobically overnight at 37 °C, and plaques were counted the following day.

### 16S rRNA gene sequencing and analysis

Amplicon sequencing data were generated and processed as previously described^38,66^. Primer sequences are listed in **Table S2**. DNA was extracted from 50 µL of culture using a DNeasy UltraClean 96 Microbial kit (Qiagen #10196-4). The V4 region of the bacterial 16S rRNA gene was amplified using Earth Microbiome Project 515F/806R primers and Accustart II PCR SuperMix (Quantabio #95137-04K) with the following thermocycler conditions: 94 °C for 3 min; 35 cycles of 94 °C for 45 s, 50 °C for 60 s, and 72 °C for 90 s; followed by 72 °C for 10 min.

PCR products were pooled in equal volumes and purified by gel extraction using a NucleoSpin Gel and PCR Cleanup Mini kit (Macherey-Nagel #740609). Libraries were prepared using a MiSeq Reagent kit v3 and sequenced with 300-bp paired-end reads on an Illumina MiSeq platform, yielding an average depth of ∼10,000 reads per sample.

Demultiplexed FASTQ files were processed using DADA2^66^. Reads were filtered and trimmed using the ‘filterAndTrim’ function with parameters truncLenF = 240, truncLenR = 160, maxEE = c(2,2), truncQ = 2, and maxN = 0. Default parameters were used for error learning and denoising. Taxonomy was assigned using the ‘assignTaxonomy’ function with the Greengenes Database (gg_13_8_train_set_97.fa). Relative abundances were calculated for each ASV, with a minimum detection threshold of 10^−3^. For visualization, undetected taxa were assigned a relative abundance of 10^−3^.

### Whole-genome sequencing of bacterial and phage isolates

Genomic DNA was extracted from 50 µL of bacterial cultures using a DNeasy UltraClean 96 Microbial kit (Qiagen #10196-4) or purified phage lysates using phenyl-chloroform extractions. DNA was quantified in 384-well plates (Bio-Rad #HSP3801, Fisher Scientific #09-761-86) using a Quant-iT PicoGreen dsDNA assay (ThermoFisher #Q33120) and normalized prior to library preparation.

Libraries were prepared using the Nextera XT DNA Library Preparation kit (Illumina) according to the manufacturer’s instructions. Briefly, DNA was subjected to tagmentation at 55 °C for 10 min, followed by neutralization and PCR amplification (10 cycles) to append Illumina sequencing adapters and sample-specific barcodes. Libraries were pooled by combining equal volumes (1 µL per sample), followed by purification using AMPure XP beads. Final library concentration was quantified using a Qubit fluorometer (ThermoFisher). Sequencing was performed on a NovaSeq X platform (25B flow cell) with 2×150 bp paired-end reads.

### Phage genome assembly and taxonomic classification

Sequencing reads from phage isolates were pre-processed by trimming, filtering, and deduplication. Genomes were assembled *de novo* using SPAdes^67^. Only contigs longer than 1 kb were retained for downstream analysis.

Pairwise genome comparisons were performed using the MUMmer^68^ alignment suite (**Table S3**). Genomes were aligned using *nucmer* with default parameters, and alignments were filtered using *delta-filter* to retain matches with ≥95% nucleotide identity. The fraction of each genome aligned at ≥95% identity was calculated using *dnadiff*. Pairwise alignments were robust to query/reference orientation. Phages were considered to belong to the same species if ≥85% of the shorter genome aligned at ≥95% nucleotide identity.

Taxonomic classification was performed using both read-based and assembly-based approaches, which yielded consistent classifications across methods. For read-based classification, filtered reads were analyzed with Phanta^69^ using default parameters and the *uhggv2_uhgv_mqplus_v1* database. Read-based classification was used to assess sample purity, with >98% of assigned reads typically belonging to a single ICTV taxon. For assembly-based classification, contigs longer than 10 kb were classified using PhaGCN as implemented in PhaBOX^70,71^. Final taxonomic assignments were reported according to the International Committee on the Taxonomy of Viruses (ICTV) taxonomy (**Table S1**).

### Anaerobic time-lapse microscopy

Time-lapse imaging of *Ef* was performed on agarose pads under anaerobic conditions. For phage treatment experiments, 1 μL of a saturated *Ef* culture diluted 1:10 in PBS was mixed with 1 µL of Tay4 (∼10^9^ PFUs/mL) and spotted onto a 1% agarose pad prepared in BHI. After drying, a coverslip was added and pads were sealed tightly with VALAP (1:1:1 vaseline:lanolin:paraffin), which maintains anaerobic conditions^72^.

For growth on spent medium, 1 μL of a saturated culture of wild-type or Δ*lpp Ef* diluted 1:100 was spotted onto 1% agarose pads composed of a 1:1 mixture of fresh BHI and spent medium. Spent medium was prepared by centrifugation of saturated cultures (8,000*g*, 10 min) in an anaerobic chamber followed by filtration through a 0.22 μm PES filter and used immediately. Agarose pads were prepared by mixing equal volumes of spent medium and molten BHI containing 2% agarose to yield a final agarose concentration of 1%. Exposure of the mixture to elevated temperature was minimized (<10 s) to limit heat-induced alterations of medium composition. Control pads were prepared using a 1:1 mixture of BHI and M9 medium (10 mL of BHI, 4 mL of 5X M9 salts (Sigma-Aldrich M9956), 6 mL of water, 40 μL of 1M MgSO_4_, 2 μL of 1M CaCl_2_, 0.2 g of agarose for 20 mL total mixture). Heat-inactivated controls were generated by heating spent medium at 95 °C for 10 min prior to pad preparation.

Phase-contrast images were acquired using a Ti-E inverted epifluorescence microscope (Nikon Instruments) equipped with a 100X oil immersion objective (numerical aperture 1.40) and a Neo 5.5 sCMOS camera (Andor Technology). Samples were maintained at 37 °C in an environmental chamber, and image acquisition was controlled using μManager v. 2.0^73^. A 5×5 field grid was imaged every 2 min for 8 h.

### Liquid chromatography-mass spectrometry (LC-MS/MS) metabolomics

Bacterial culture supernatants were collected by centrifugation (4,000*g*, 10 min, 4 °C) and immediately frozen at –70 °C. Samples were prepared as previously described^47^. Samples were thawed once immediately prior to analysis, maintained on ice, and homogenized by pipetting. Aliquots (20 µL) were transferred to a 96-well polypropylene plate kept on ice, and pooled quality control (QC) samples were generated by combining 5 μL from each sample and processed in parallel. QC samples were injected at regular intervals throughout the run to monitor instrument performance. Procedural blanks (water in place of sample) were included to assess background signal. Sample injection order was randomized to minimize acquisition bias.

Metabolites were extracted by adding 80 μL of a pre-chilled (–20 °C) acetonitrile:methanol (1:1) solution containing 5% water and stable isotope-labelled internal standards. Plates were equilibrated at –20 °C for 1 h to precipitate proteins, followed by centrifugation (6,000*g*, 5 min, –9 °C). Supernatants were transferred to a new plate, and 2 µL were injected for analysis. Internal standards were used to monitor extraction efficiency and chromatographic consistency.

Samples were analyzed by hydrophilic interaction chromatography (HILIC) coupled to a Thermo Q Exactive HF Orbitrap mass spectrometer using electrospray ionization in both positive and negative-modes (separate injections). Chromatographic^74^ and mass spectrometric parameters^75^ were as previously described with minor modifications. Data were acquired in full MS-ddMS2 mode over a mass range of 60-900 *m*/*z*, with resolutions of 60,000 (MS1) and 15,000 (MS2), loop count of 4, and an isolation window of 1.2 Da. An inclusion list was used to prioritize MS2 acquisition of metabolites in an in-house library, with additional data-dependent MS2 collection.

### Metabolomics data analysis

Metabolomics data were processed and analyzed as previously described^47^. Raw data were processed using MS-DIAL v. 4.60^76,77^ with an MS1 tolerance of 0.01 Da, MS2 tolerance of 0.015 Da, and a minimum peak height of 100,000 detected in at least two of three biological replicates. Peak alignment was performed with a retention time tolerance of 0.05 min and mass tolerance of 0.015 Da.

Metabolite annotations were assigned based on MS2 spectra using an in-house library of standards analyzed under identical chromatographic conditions, supplemented with the MassBank of North America MS2 repository (https://mona.fiehnlab.ucdavis.edu/) for metabolites not present in the internal library. Features were retained for downstream analysis if they were detected in at least two of three biological replicates and exceeded the water blank average signal by >5-fold in at least one sample, ensuring robust detection above background.

Metabolites were defined as depleted if their peak areas decreased by >100-fold relative to fresh BHI medium, had a peak intensity >5,000 in BHI, and exhibited statistically significant differences (two-sample Student’s *t*-test with Benjamini-Hochberg correction, adjusted *p*<0.05). For fold-change calculations, zero values were set to 1.

Niche overlap between species was estimated by comparing the shared set of depleted features between species grown in isolation in BHI relative to the total number of features depleted by each species.

### Single-cell liquid isolation of phage-resistant *Ef* mutants

To isolate Tay4-resistant *Ef* strains, *Ef* was grown with Tay4 (MOI∼1) either in a monoculture or within the MD2 SIC in BHI supplemented with 1 mM CaCl_2_ for 48 h at 37 °C with continuous shaking. Because maintenance of phage resistance required continuous phage pressure, single-cell isolation was performed in liquid culture to avoid phage contamination risks associated with spreading lysate on plates.

For isolation from monocultures, saturated cultures (10^9^ CFUs/mL) were diluted 10^-9^ into 96-well plates containing fresh BHI with or without Tay4 and incubated anaerobically for 48 h at 37 °C with continuous shaking. Under these conditions, most wells (∼90%) exhibited no growth, indicating limiting dilution to approximately single-cell inoculation (**Fig. S4e**). Comparable growth frequencies and lag times were observed in the presence and absence of Tay4, consistent with isolation of phage-resistant cells when phage was included.

For isolation from communities, saturated cultures (10^8^ *Ef* CFUs/mL) were diluted 10^-8^ into 96-well plates containing BHI supplemented with Tay4 and incubated anaerobically for 48 h at 37 °C with continuous shaking. Wells containing *Ef* were identified based on growth dynamics, colony morphology after aerobic plating, and single-cell imaging.

Isolates were re-passaged for 48 h under anaerobic conditions (37 °C, continuous shaking) with and without Tay4 to confirm phage resistance. All isolates were subjected to whole-genome sequencing to verify strain identity and assess sample purity. Cultures were then frozen at –70 °C for downstream analyses.

### Long-read *Ef* whole genome sequencing and assembly

Saturated cultures were pelleted by centrifugation (4000*g*, 10 min). To preserve high-molecular-weight DNA for closed genome assembly, pellets were resuspended in lysis buffer (50 mM Tris-HCl, 1 mM EDTA, 0.5% SDS, 200 µg/mL proteinase K) and incubated at 37 °C for 1 h.

DNA was extracted using phenyl-chloroform as described above, or by resuspension of pellets in DNA/RNA Shield (Zymo Research #R1100) followed by extraction by Plasmidsaurus. Long-read sequencing and *de novo* assembly were performed by Plasmidsaurus using Oxford Nanopore Technologies platforms.

### Identification of genetic variants associated with phage resistance

To identify genetic changes associated with Tay4 resistance, short-read sequencing data from *Ef* isolates were analyzed for single nucleotide polymorphisms (SNPs) and structural variants. SNP calling was performed using Clair3^78^ with a model trained on Illumina paired-end reads. Variants were called using the following parameters: *no_phasing_for_fa*, *include_all_ctgs*, *haploid_sensitive*, *enable_long_indel*, and *enable_variant_calling_at_sequence_head_and_tail*, with additional filtering requiring a minimum coverage of 5 reads and a minimum mapping quality of 10.

To facilitate visualization of mutation patterns across the genome, SNPs separated by less than 1 kb were grouped into clusters. This analysis did not reveal consistent SNP patterns across isolates, motivating further interrogation of structural variation.

Structural variants, including deletions, duplications, insertions, and translocations, were identified using DELLY^79^ with default parameters, except that the minimum number of supporting reads was reduced to 5 to increase sensitivity to low-frequency variants. Genome-wide structural variation and rearrangements were further assessed by generating synteny plots from alignments of assembled genomes of resistant isolates to the wild-type *Ef* genome using custom Python scripts.

### Quantification of *hyx* inverton orientation

To quantify the orientation of the promoter region upstream of the *hyx* operon during phage treatment, *Ef* was grown with Tay4 (MOI∼1) in monoculture or within the MD2 SIC in BHI supplemented with 1 mM CaCl_2_ for 48 h at 37 °C without shaking. Cultures were passaged 1:200 into fresh BHI+CaCl_2_ for one additional passage, and samples were collected at 0, 4, 8, 24, and 48 h for each passage. Cells were pelleted (4000*g*, 10 min), washed twice with PBS to remove extracellular DNA, resuspended in PBS, and stored at –70 °C prior to processing. DNA was extracted from 50 µL of each sample using a DNeasy UltraClean 96 Microbial kit (Qiagen #10196-4).

The inverton region was amplified using a two-step PCR approach with custom primers (**Table S2**). The first PCR was performed using Accustart II PCR SuperMix (Quantabio #95137-04K) with the following conditions: 94 °C for 3 min; 35 cycles of 94 °C for 45 s, 52 °C for 60 s, and 72 °C for 30 s; followed by 72 °C for 10 min. Products were diluted 1:500 and used as input for a second PCR to append sequencing adapters and indices (**Table S2**) with the following conditions: 94 °C for 3 min; 15 cycles of 94 °C for 45 s, 54 °C for 60 s, and 72 °C for 90 s; followed by 72 °C for 10 min.

Amplicons were pooled in equal volumes and purified by gel extraction with a NucleoSpin Gel and PCR Cleanup Mini kit (Macherey-Nagel #740609). Libraries were prepared using an Illumina MiSeq Reagent kit v3 and sequenced with 300-bp paired-end reads, yielding an average depth of ∼40,000 reads per sample. Demultiplexed FASTQ files were processed using *cutadapt*^80^ for adapter trimming (default parameters) and *bowtie2*^81^ for alignment to the reference sequence, excluding unmapped reads. High alignment rates (>99% of reads per sample) indicated efficient and specific amplification of the target region.

### Re-derivation of stool-derived communities under phage treatment

Frozen stool samples previously used to derive SD1, SD2, MD1, and MD2^38^ were thawed, and 2 µL of each sample were inoculated 1:100 into 200 µL of BHI supplemented with 1 mM CaCl_2_, with or without addition of 1 µL of the phage cocktail. Cultures were grown anaerobically for 48 h at 37 °C without shaking and subsequently passaged 1:200 for six additional passages in fresh BHI without phage to allow community recovery. After the seventh passage, communities were aliquoted into 25% glycerol stocks and samples for downstream analyses and frozen at –70 °C.

### *Salmonella* invasion and reintroduction assays

Complete and Δ*Ef* SICs were revived in BHI and passaged three times (1:200, 48 h per passage) at 37 °C under anaerobic, non-shaking conditions to establish steady-state communities prior to invasion. *Salmonella enterica* serovar Typhimurium SL1344 (*S*Tm) was streaked from a frozen stock onto BHI agar (1.5%) and grown anaerobically at 37 °C for 24 h. A single colony was then inoculated into BHI and cultured for 48 h under the same conditions to match SIC passaging.

For invasion assays, SICs were mixed with *S*Tm at defined ratios by combining 1 µL of SIC with 1 µL of saturated, 1:10 diluted, or 1:100 diluted *S*Tm, corresponding to initial *S*Tm fractions of 50%, 10%, and 1%, respectively. Co-cultures were passaged four times (1:200 dilution, 48 h per passage) in BHI without shaking.

To assess the ability of *Ef* to displace established *S*Tm, *Ef* was reintroduced into Δ*Ef* SICs after *S*Tm invasion. Co-cultures containing 10% initial *S*Tm were mixed with wild-type *Ef* at 50%, 10%, or 1% initial fractions using saturated or diluted cultures as above, and passaged for four additional passages under the same conditions.

Samples were collected after each passage and stored at –70 °C for downstream analyses. *S*Tm abundance was quantified by plating serial dilutions onto LB agar supplemented with 50 µg/mL streptomycin, which selectively supports *S*Tm growth. Plates were incubated aerobically at 37 °C overnight, providing an additional selection for *S*Tm, and colonies were counted the following day. Quantification was performed after passages 1, 4, 5, and 8.

### Construction of a barcoded transposon mutant library in *Ef*

A barcoded transposon insertion library was generated by conjugation between *E. coli* donor cells carrying the vector PHLL250_NN1 and *Ef* recipient cells at a 1:1 ratio. Donor cells (AMD290; WM3064+pHLL250_NN1) were grown to exponential phase in LB Lennox medium (BD #240230) supplemented with 50 μg/mL carbenicillin and 300 μM diaminopimelic acid (DAP) at 37 °C, harvested, washed, and resuspended in LB+DAP to OD_600_∼40. *Ef* recipient cells from an overnight culture were similarly prepared and resuspended to OD_600_∼40.

Equal volumes (100 µL each) of donor and recipient suspensions were mixed and spotted onto a 0.45 μm membrane filter placed on LB+DAP agar plates. After overnight incubation at room temperature, cells were recovered, resuspended in LB, and plated on LB agar containing 40 μg/mL kanamycin to select for transposon insertions. Colonies appeared within 24 h of aerobic incubation at 37 °C.

Colonies were pooled and expanded aerobically in LB supplemented with 40 μg/mL kanamycin to generate the mutant library. Saturated cultures were aliquoted into single-use glycerol stocks (15% final glycerol) and stored for downstream applications. Genomic DNA was extracted (DNeasy Blood and Tissue DNA Kit, Qiagen) for TnSeq-based mapping of insertion sites and association with DNA barcodes. The final mutant library contains more than one million uniquely barcoded insertion mutants, providing near saturating coverage of the *Ef* genome.

### Mapping transposon insertion sites to barcodes by TnSeq

Transposon insertion sites and associated DNA barcodes in the *Ef* library were identified using an updated TnSeq protocol based on a previous method^59^. Key modifications included the use of two rounds of PCR to amplify insertion junctions and a splinkerette adapter design in place of a Y-adapter, improving recovery of insertion-barcode associations.

Briefly, 1 µg of genomic DNA from the barcoded transposon library was sheared to ∼300 bp using a Covaris instrument. DNA end repair, A-tailing, and ligation of splinkerette adapters were performed using a KAPA Hyper Prep kit (Roche #KK8504) according to the manufacturer’s instructions. Splinkerette adapters were prepared by annealing oligonucleotides (Adapt_1_tnseqV3 and Adapt_2_tnseqV3; **Table S2**) using a controlled temperature ramp from 97.5 °C to 4 °C. Adapter-ligated DNA was purified by bead cleanup and used as input for a first round of PCR with primers targeting the transposon–genome junction (Rd1_FOR_tnseqV3, Rd1_REV_tnseqV3; **Table S2**) using KAPA HiFi HotStart ReadyMix (Roche #KK2602).

The first PCR consisted of an initial denaturation at 98 °C for 45 s, followed by 15 cycles of 98 °C for 15 s, 60 °C for 30 s, and 72 °C for 30 s, and a final extension at 72 °C for 1 min. Products were purified by double-sided bead cleanup and used as input for a second PCR to append Illumina sequencing adapters and dual indices, enabling multiplexing and identification of index hopping events. The second PCR was performed with an initial denaturation at 98 °C for 45 s, followed by 8 cycles of 98 °C for 15 s, 60 °C for 30 s, and 72 °C for 30 s, and a final extension at 72 °C for 1 min.

Final libraries were purified by double-sided bead cleanup and assessed for quality using a High Sensitivity D1000 ScreenTape assay (Agilent 4200 TapeStation). Libraries were sequenced on an Illumina NovaSeq platform (Novogene). To ensure comprehensive mapping of insertion sites, three independent libraries were generated and sequenced. Sequencing data were processed using previously established pipelines^59^ to map insertion sites, associate barcodes, and define the set of mutants suitable for downstream fitness analyses.

### Transposon mutant library competition assays

Two glycerol stocks of the *Ef* transposon library were thawed and inoculated into 50 mL of BHI or BHI supplemented with 1 mM CaCl_2_ (for phage treatment) and grown anaerobically for 48 h at 37 °C without shaking. To maintain representation of library diversity, a minimum input of 10 µL of saturated culture was used for each experiment, corresponding to an estimated ≥10X coverage for each barcode. The library was grown either in monoculture or co-cultured with SICs by combining 10 µL of the saturated library with 30 µL of SIC and inoculating into 4 mL of BHI. Cultures were incubated anaerobically for 48 h at 37 °C without shaking. After growth, aliquots (250 and 50 µL) were collected, pelleted (4,000*g*, 10 min), and stored at –70 °C for BarSeq and 16S rRNA sequencing, respectively.

### Measurement of mutant fitness by barcode sequencing (BarSeq)

Genomic DNA from all BarSeq samples was extracted in a 96-well format using a QIAamp 96 DNA QIAcube HT kit (Qiagen #51331) according to the manufacturer’s instructions, with the exception that a vacuum manifold was used for sample processing. BarSeq libraries were prepared as previously described^59^ using unique dual-indexed primers (**Table S4**) to enable multiplexing and minimize index misassignment.

Libraries were sequenced on an Illumina NovaSeq X 10B platform (1×100 bp). Mutant fitness was quantified by normalizing barcode abundances in each condition to their corresponding abundances at the initial time point (Time 0), and comparing fold changes between community and monoculture growth conditions.

### Construction of gene knockout strains in *Ef*

Gene knockouts were generated using λ Red recombineering^82^ under aerobic conditions. Strains, primers, and plasmids are listed in **Tables S1** and **S2**. PCR products containing a chloramphenicol resistance cassette (cm^R^) flanked by ∼50 bp homology arms targeting the *Ef* genome were amplified from pKD3 in 50 µL reactions (0.5 µL of Phusion High-Fidelity DNA polymerase, 10 µL of 5X HF buffer, 1 µL of 10 mM dNTPs, 2.5 µL of 10 µM forward primer, 2.5 µL of 10 µM reverse primer, 1.5 µL of DMSO, 5 µL of pKD3 template, with remaining water; NEB #M0530). PCR used the following conditions: 98 °C for 30 s; 35 cycles of 98 °C for 10 s, 62 °C for 30 s, and 72 °C for 30 s; and 72 °C for 10 min. Products were verified by 1% agarose gel electrophoresis and Sanger sequencing prior to purification (Qiagen PCR cleanup kit, #28104).

The 11 Red recombinase plasmid pSIM6 was electroporated into 50 µL of freshly prepared electrocompetent *Ef* cells (2.5 kV, 25 mF, 200 Ω), and transformants were selected on LB-agar plates supplemented with 100 µg/mL ampicillin at 30 °C. Cells carrying pSIM6 were grown to mid-log phase (OD_600_ 0.4–0.6) at 30 °C, heat-induced at 42 °C for 15 min to activate recombinase expression, and rapidly cooled on ice. Cells were washed with ice-cold water twice and electroporated (2.5 kV, 25 mF, 200 Ω) with PCR amplicons containing the cm^R^ cassette and homology arms. Recombinants were selected on LB-agar plates supplemented with 20 µg/mL chloramphenicol and purified by repeated streaking. Loss of the pSIM6 plasmid was confirmed by absence of growth on 100 µg/mL ampicillin-containing plates.

Correct insertion of the cm^R^ at the target locus was verified by colony PCR and Sanger sequencing. The cm^R^ cassette was subsequently excised using FLP recombinase by introducing the temperature-sensitive plasmid pCP20 via electroporation as above. Transformants were selected at 30 °C 100 µg/mL ampicillin-containing plates and then re-struck at 42 °C on non-selective media to cure pCP20. Successful excision of cm^R^ and loss of pCP20 were confirmed by antibiotic sensitivity and colony PCR. Final knockout strains were verified by PCR and Sanger sequencing and stored in 25% glycerol at –70°C.

## Data and code availability

Custom code used for data analysis with corresponding non-sequencing data is available at https://doi.org/10.5281/zenodo.20057640. Additional information required to reproduce or reanalyze the results is available from the corresponding author upon reasonable request.

## Acknowledgements

We thank members of the Huang lab for helpful discussions. This work was funded by an NSF Graduate Research Fellowship and a Siebel Scholars Fellowship (to T.H.N.), a James S. McDonnell Postdoctoral Fellowship (to H.S.), a Stanford Propel Postdoctoral Scholars Fellowship (to S.M.-M), an HHMI Hanna G. Gray Fellowship (to S.M.-M), NSF Award EF-2125383 (to K.C.H.), and NIH Awards R35 GM150996 (to A.J.H.), RM1 GM135102 (to A.M.D. and K.C.H.), and DP1 DK147449 (to K.C.H.). K.C.H. is a Chan Zuckerberg Biohub Investigator. Some of the computing for this project was performed on the Sherlock cluster. We thank Stanford University and the Stanford Research Computing Center for providing computational resources and support that contributed to these research results.

## Author Contributions Statement

Conceptualization, T.H.N. and K.C.H..; methodology, T.H.N., M.S., V.T., J.S., H.S., A.J.H.; investigation, T.H.N., M.S., N.T.L., V.T., J.S., H.S., S.M.-M., J.A.L., P.-Y.H.;

writing—original draft, T.H.N., K.C.H..; writing—review & editing, all authors; funding acquisition, A.M.D., A.J.H., K.C.H.; resources, A.M.D., A.J.H., K.C.H.; supervision, A.M.D., A.J.H., K.C.H.

