## Supplementary information for "Phage-based microbiome manipulation reveals ecological interactions within gut communities"

**Supplementary information and data: Phage-based microbiome manipulation reveals ecological interactions within gut communities**

**Authors:**

Taylor H. Nguyen[1], Morgan Su[2], Nhien T. Lu[1], Valentine Trotter[3], Saria A. McKeithen-Mead[1,2], Jamie Alcira Lopez[1,4], Jiawei Sun[1], Zachary Hallberg[5], Handuo Shi[1,2], Po-Yi Ho[1,6], Brian C. DeFelice[7], Michiko E. Taga[5], Adam M. Deutschbauer[3,5], Andrew J. Hryckowian[8,9], Kerwyn Casey Huang[1,2,7,†]

**Affiliations:**

[1]Department of Bioengineering, Stanford University, Stanford, CA 94305

[2]Department of Microbiology and Immunology, Stanford University School of Medicine, Stanford, CA 94305

[3]Environmental Genomics and Systems Biology Division, Lawrence Berkeley National Laboratory, Berkeley, CA 94720, USA

[4]Department of Applied Physics, Stanford University, Stanford, CA 94305

[5]Department of Plant and Microbial Biology, University of California, Berkeley, CA 94720, USA

[6]Center for Interdisciplinary Studies, Westlake University, Hangzhou, China

[7]Chan Zuckerberg Biohub, San Francisco, CA 94158

[8]Department of Medicine, Division of Gastroenterology and Hepatology, University of Wisconsin School of Medicine and Public Health, Madison, WI 53706, USA

[9]Department of Medical Microbiology & Immunology, University of Wisconsin School of Medicine and Public Health, Madison, WI 53706, USA

**Supplemental Tables**

**Table S1:** Bacterial strains, communities, and phages used in this study.

**Table S2:** Oligonucleotides and plasmids used in this study.

**Table S3:** Genome alignment and sequence similarity of phages used in the phage cocktail.

**Table S4:** Barseq primers used for sequencing the *Ef* transposon mutant library in fitness assays.

**Supplemental Videos**

**Movie S1:** Time-lapse microscopy of *Ef* treated with Tay4 (MOI~1) under anaerobic conditions on a 1% agarose pad containing BHI. Twenty-five fields of view were collected. Representative movie shown. Scale bar: 5 µm.

**Movie S2:** Time-lapse microscopy of *Ef* Δ*lpp* grown on a 1% agarose pad containing 50% fresh BHI and 50% M9 salts (**Methods**; **Fig. 6e**). Cells exhibit normal growth. Data were collected on two independent days with 25 fields of view per day (~1-10 initial cells per field). Representative movie shown. Scale bar: 5 µm.

**Movie S3:** Time-lapse microscopy of wild-type *Ef* grown on a 1% agarose pad containing 50% fresh BHI and 50% M9 salts (**Methods**; **Fig. S11a**). Cells exhibit normal growth. Data were collected on two independent days with 25 fields of view per day (~1-10 initial cells per field). Representative movie shown. Scale bar: 5 µm.

**Movie S4:** Time-lapse microscopy of *Ef* Δ*lpp* grown on a 1% agarose pad containing 50% fresh BHI and 50% MD2-Δ*Ef* spent medium (**Methods**; **Fig. 6e**). Cells exhibit swelling, elongation, and lysis. Data were collected on two independent days with 25 fields of view per day (~1-10 initial cells per field). Representative movie shown. Scale bar: 5 µm.

**Movie S5:** Time-lapse microscopy of wild-type *Ef* grown on a 1% agarose pad containing 50% BHI and 50% MD2-Δ*Ef* spent medium (**Methods**; **Fig. S11b**). No morphological changes are observed under these conditions. Data were collected on two independent days with 25 fields of view per day (~1-10 initial cells per field). Representative movie shown. Scale bar: 5 µm.

**Movie S6**: Time-lapse microscopy of *Ef* Δ*lpp* grown on a 1% agarose pad containing 50% fresh BHI and 50% heat-treated MD2-Δ*Ef* spent medium (95 °C, 15 min; **Methods**; **Fig. S11c**). Cells exhibit normal growth under these conditions, indicating loss of the active factor driving antagonism upon heat treatment. Data were collected on two independent days with 25 fields of view per day (~1-10 initial cells per field). Representative movie shown. Scale bar: 5 µm.

**Movie S7**: Time-lapse microscopy of *Ef* Δ*lpp* grown on a 1% agarose pad containing 50% PBS and 50% MD2-Δ*Ef* spent medium (**Methods**; **Fig. S11d**). Cells exhibit normal morphology under conditions of reduced growth. Data were collected on two independent days with 25 fields of view per day (~1-10 initial cells per field). Representative movie shown. Scale bar: 5 µm.

**Movie S8**: Time-lapse microscopy of *Ef* Δ*lpp* grown on a 1% agarose pad containing 50% fresh BHI and 50% spent medium from the 14-member synthetic community (**Methods**; **Fig. S11f**). Cells exhibit normal growth under these conditions, suggesting that other members present in the complete MD2 community contribute to the full antagonistic phenotype. Data were collected on two independent days with 25 fields of view per day (~1-10 initial cells per field). Representative movie shown. Scale bar: 5 µm.

### Supplementary Figures

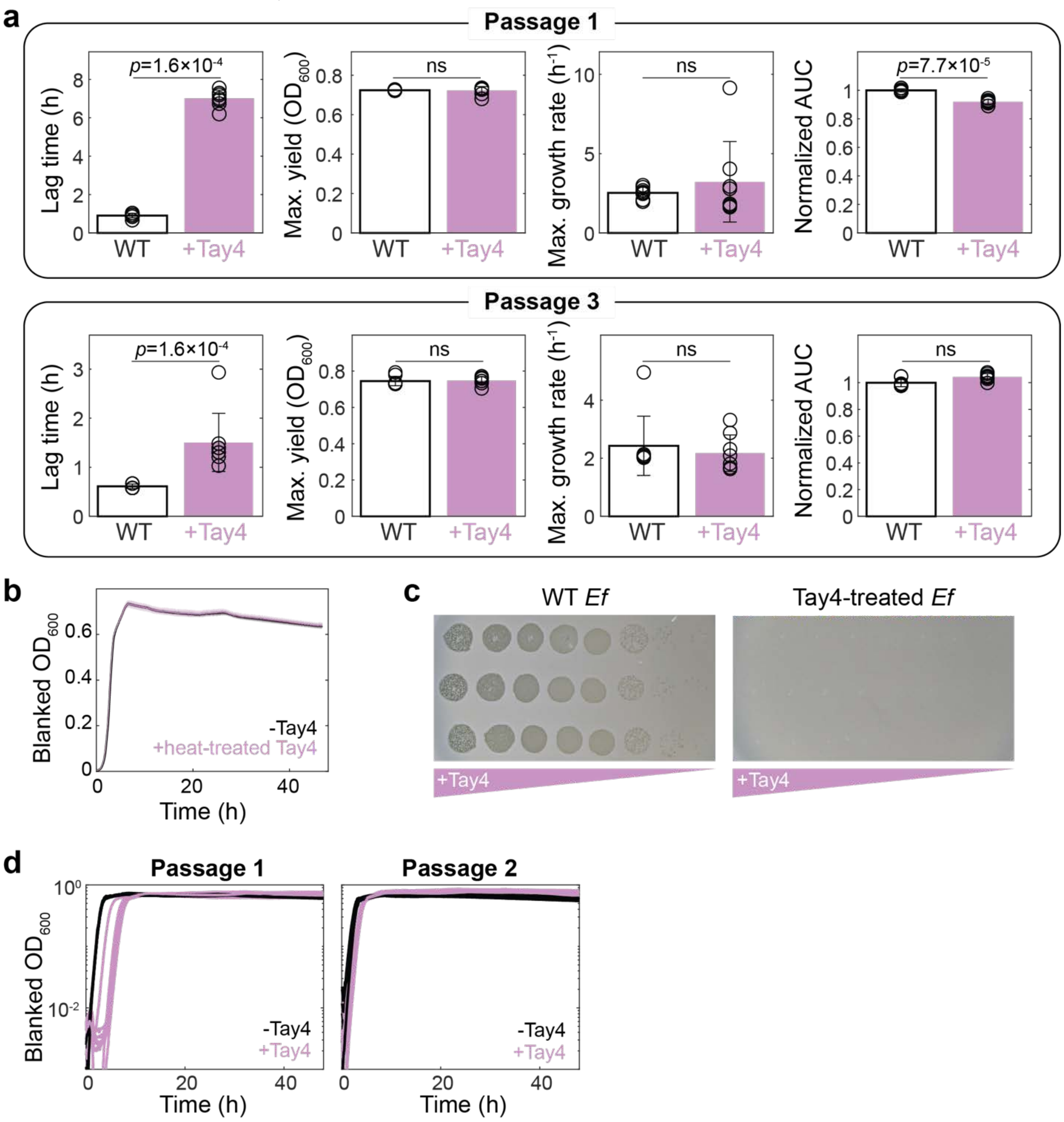


**Figure S1: Phage treatment does not reduce *Ef* fitness in monoculture.**

a) Growth metrics of *Ef* following Tay4 treatment in passage 1 and passage 3 (**Fig. 1b**). Bars indicate mean±s.d. ($n$=8 biological replicates), with individual replicates overlaid. Lag time was defined as the time to reach $OD_{600}$=0.01. Maximum growth rate was calculated as the maximum derivative of ln($OD_{600}$) after smoothing. AUC (0–48 h) was normalized to the mean AUC of untreated wild-type (WT) *Ef*.

b) Growth dynamics of *Ef* in the presence of heat-denatured Tay4 (95 °C, 10 min). Background-subtracted $OD_{600}$ curves are shown for cultures with (purple) or without (black) heat-treated phage. No lag increase was observed, indicating that growth delays require active phage. Thus, for all experiments hereafter we compare phage-treated cultures to untreated cultures as controls. Curves show mean±s.d. (*n*=3 biological replicates).

c) Plaque assays of Tay4 on wild-type (WT) *Ef* and previously Tay4-treated *Ef*, showing loss of susceptibility in phage-exposed populations.

d) Growth dynamics of *Ef* during two consecutive passages with Tay4 (MOI~1; purple) compared to untreated *Ef* (black). Background-subtracted $OD_{600}$ curves show recovery of growth kinetics across passages despite continued phage pressure. Each curve represents a biological replicate (*n*=8).

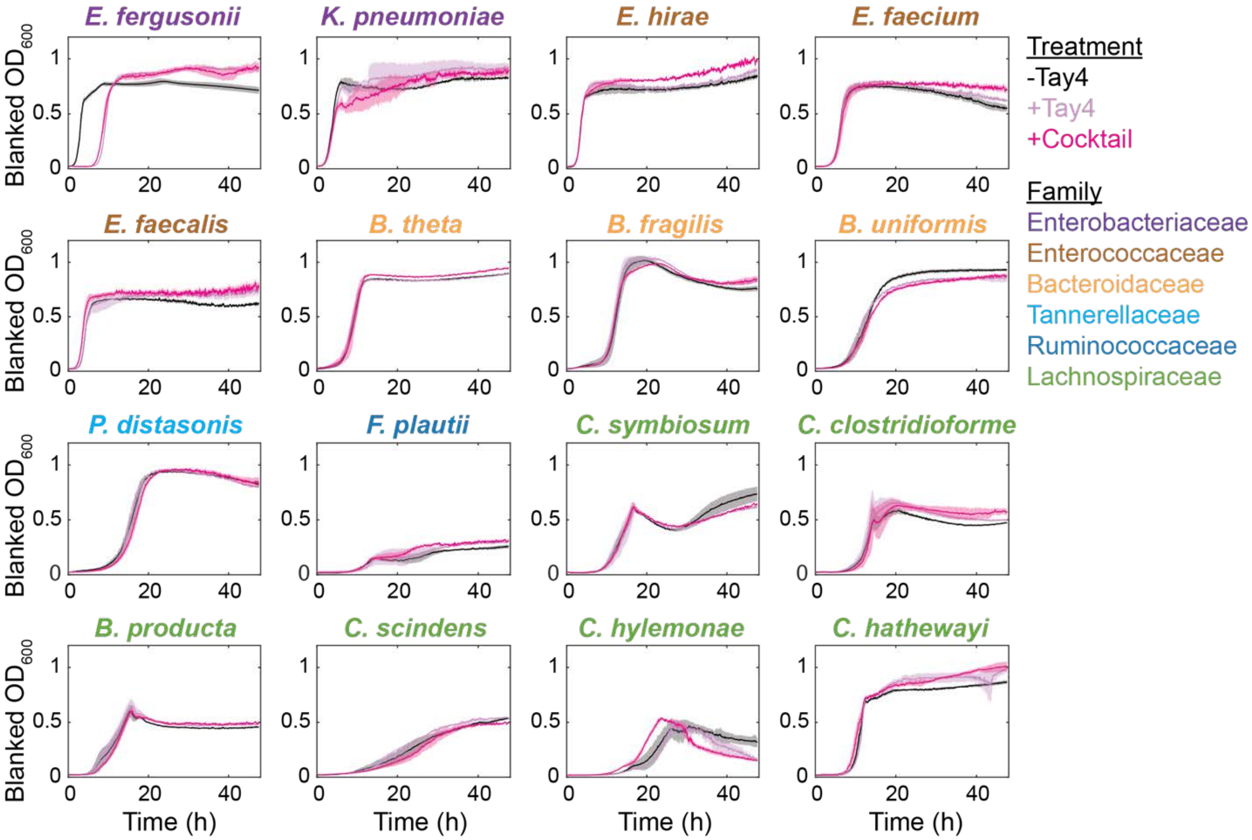


**Figure S2: Tay4- and cocktail-mediated lysis is specific to *Ef* within a 15-member community.**

Background-subtracted $OD_{600}$ growth dynamics are shown for each species grown without phage (black), with Tay4 (purple; MOI~1), or with a phage cocktail (magenta; MOI~1). Lines represent mean±s.d. across biological replicates (*n*=3). Species names are colored by taxonomic family. Tay4 treatment selectively inhibits growth of *Ef*, with no detectable growth defects observed in other community members.

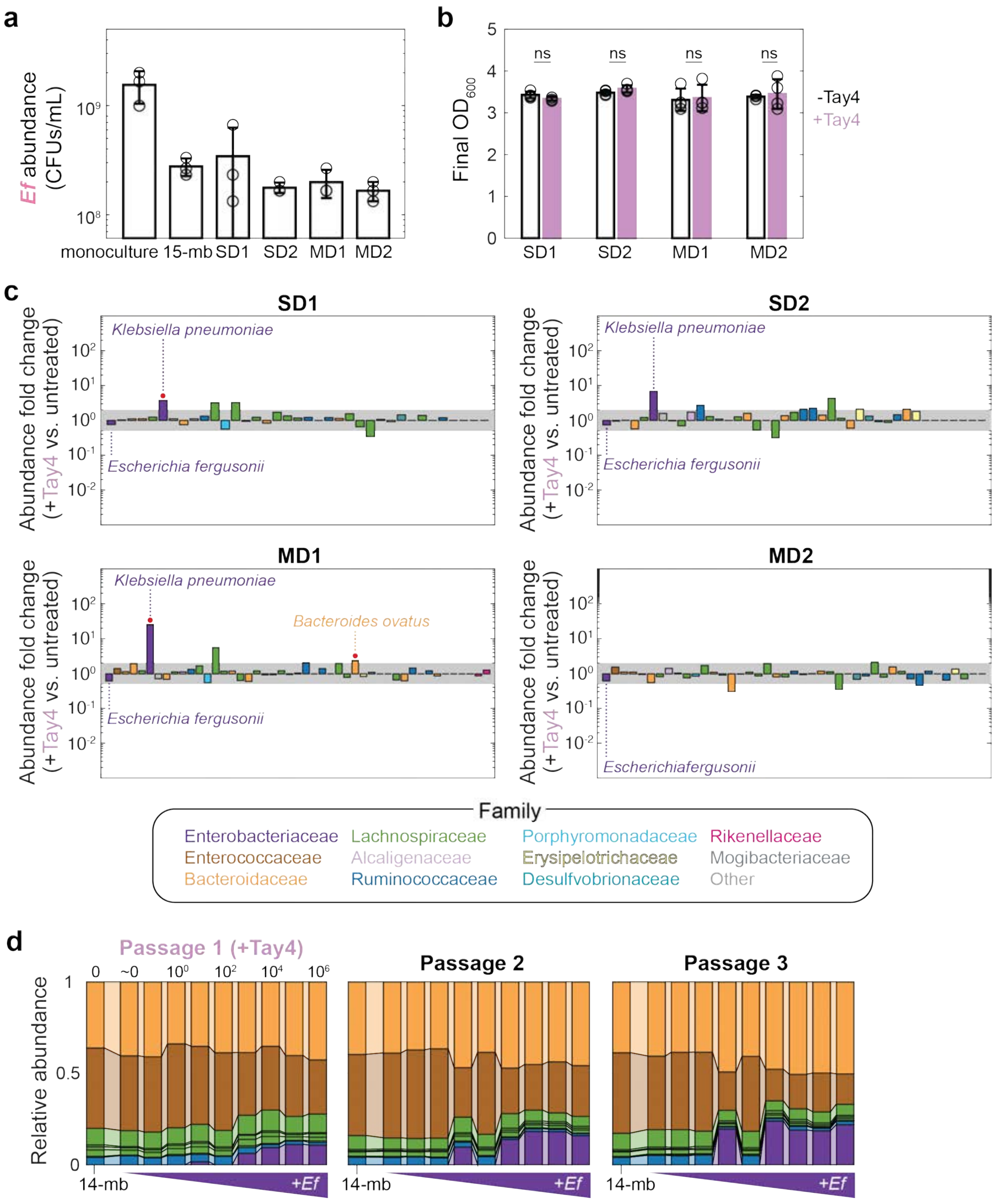


**Figure S3: Phage treatment reduces *Ef* abundance without altering overall community yield.**

a) Absolute abundance of *Ef* (CFUs/mL) in monoculture, a 15-member (15-mb) synthetic community, and four complex SICs (SD1, SD2, MD1, and MD2). *Ef* was quantified by selective plating and colony morphology (**Methods**). Measurements were taken after ≥4 passages to ensure steady-state composition. Bars show mean±s.d. ($n$=3 biological replicates), with individual replicates overlaid.

b) Final community biomass following Tay4 treatment. $OD_{600}$ is shown for communities grown with (purple; MOI~1) or without (white) Tay4. Differences were not significant (ns; two-sided Student's $t$-test; $p$-values>0.05). Bars show mean±s.d. ($n$=3 biological replicates) with individual replicates overlaid.

c) Community composition changes following Tay4 treatment. Fold change in ASV abundance is shown for Tay4-treated versus untreated communities. Bars represent mean across biological replicates ($n$=3). The shaded region indicates a 2-fold increase/decrease. ASVs outside this region with significant changes (two-sample Student's $t$-test with Benjamini-Hochberg correction, adjusted $p$<0.05) are marked with asterisks. ASVs that newly emerge above or drop below the limit of detection ($10^{-3}$) are highlighted (red circles). Colors denote taxonomic family.

d) Synthetic community composition following *Ef* titration and Tay4 treatment. A 14-member community (14-mb; no *Ef*) and 15-member communities with varying Ef inputs were treated with Tay4 in passage 1 and passaged for two recovery passages (1:200 dilution). Community composition remained largely stable despite reduced *Ef* abundance. ASVs are colored by family.

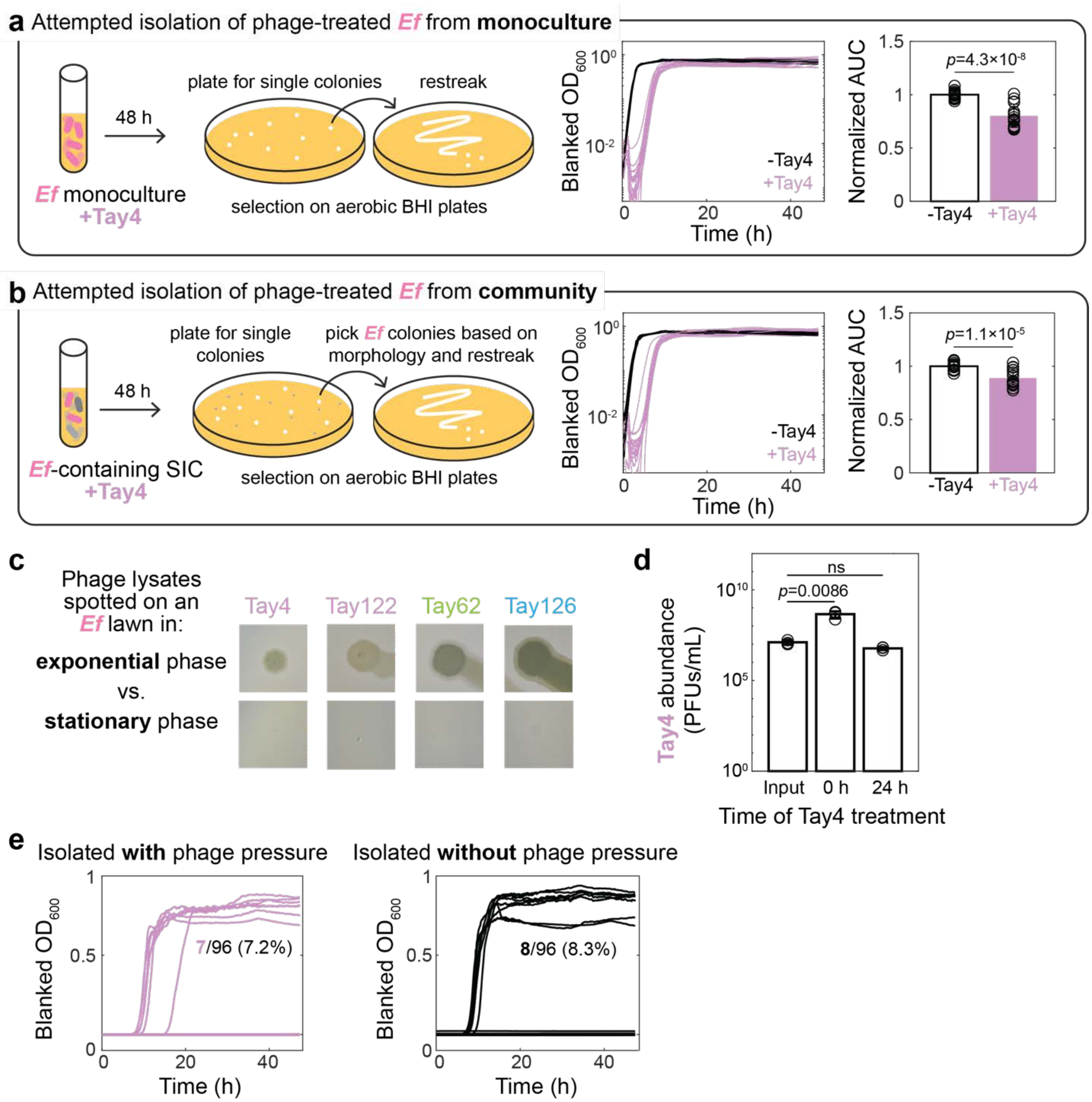


**Figure S4: *Ef* resistance to Tay4 requires continual phage pressure and phage lytic activity is reduced in stationary phase.**

a) Schematic of phage-treated *Ef* isolation from monoculture using plates. Cultures grown with Tay4 (MOI~1) were plated for single colonies, re-streaked, and regrown before testing for resistance. Growth dynamics (left) and normalized AUC (right, 0–48 h) show that isolates lost resistance upon passage without phage. Curves represent biological replicates (*n*=8).

b) Schematic of phage-treated *Ef* isolation from community using plates. The *Ef*-containing MD2 SIC was treated with Tay4 (MOI~1) for 48 h, plated, and *Ef* colonies were isolated based on morphology before testing for resistance. As in monoculture, growth dynamics (left) and normalized AUC (right, 0–48 h) show that isolates lost resistance after removal of phage pressure. Curves represent biological replicates (*n*=8).

c) Growth-phase dependence of phage lysis. Plaque assays were performed on lawns of *Ef* in exponential or stationary phase. Phages formed plaques on exponentially growing cells but not on stationary-phase cells, indicating reduced lytic activity in stationary phase.

d) Limited propagation of Tay4 in stationary-phase cultures. Tay4 titers are shown following addition to *Ef* cultures at 0 h or 24 h of growth. Phage amplification is reduced when added to stationary-phase cultures. Bars shown mean±s.d. (*n*=3 biological replicates) with individual replicates overlaid.

e) To test whether *Ef* reverted to a susceptible state after one passage without phage pressure, plates were grown with (left) and without (right) Tay4 (**Methods**). Only 7 or 8 wells in each 96-well plate exhibited growth, indicating that dilution led to inoculation of either 0 or 1 *Ef* cell in each well.

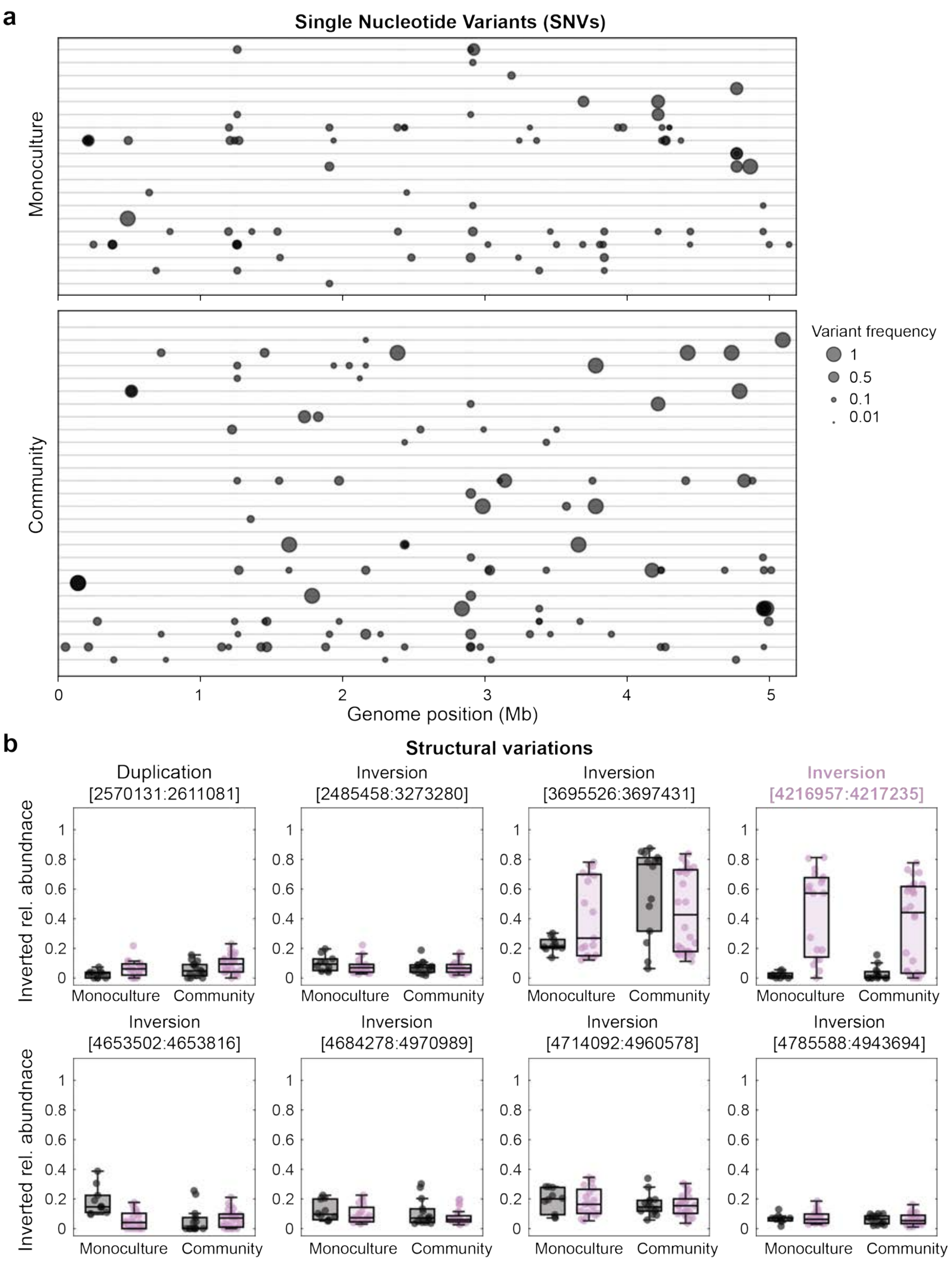


**Figure S5: Phage resistance is not associated with consistent single nucleotide variants (SNVs) but is linked to structural variation.**

a) Genome-wide distribution of SNVs in *Ef* isolates following Tay4 treatment in monoculture (top) and the MD2 community (bottom). Each point represents a SNV, positioned by genomic coordinate and scaled by variant frequency. No consistent or recurrent SNVs were observed across isolates in either condition.
b) Structural variation analysis of *Ef* isolates. Relative abundance of structural variants (duplications and inversions) is shown for monoculture (black) and MD2 community (purple) isolates. While most structural variants occur at low frequency, a subset of inversions were enriched during phage treatment. The inversion upstream of the *hyx* operon is the only variant that was specific to phage treatment, consistent with a role in resistance. Box plots show median, interquartile range, and individual replicates.

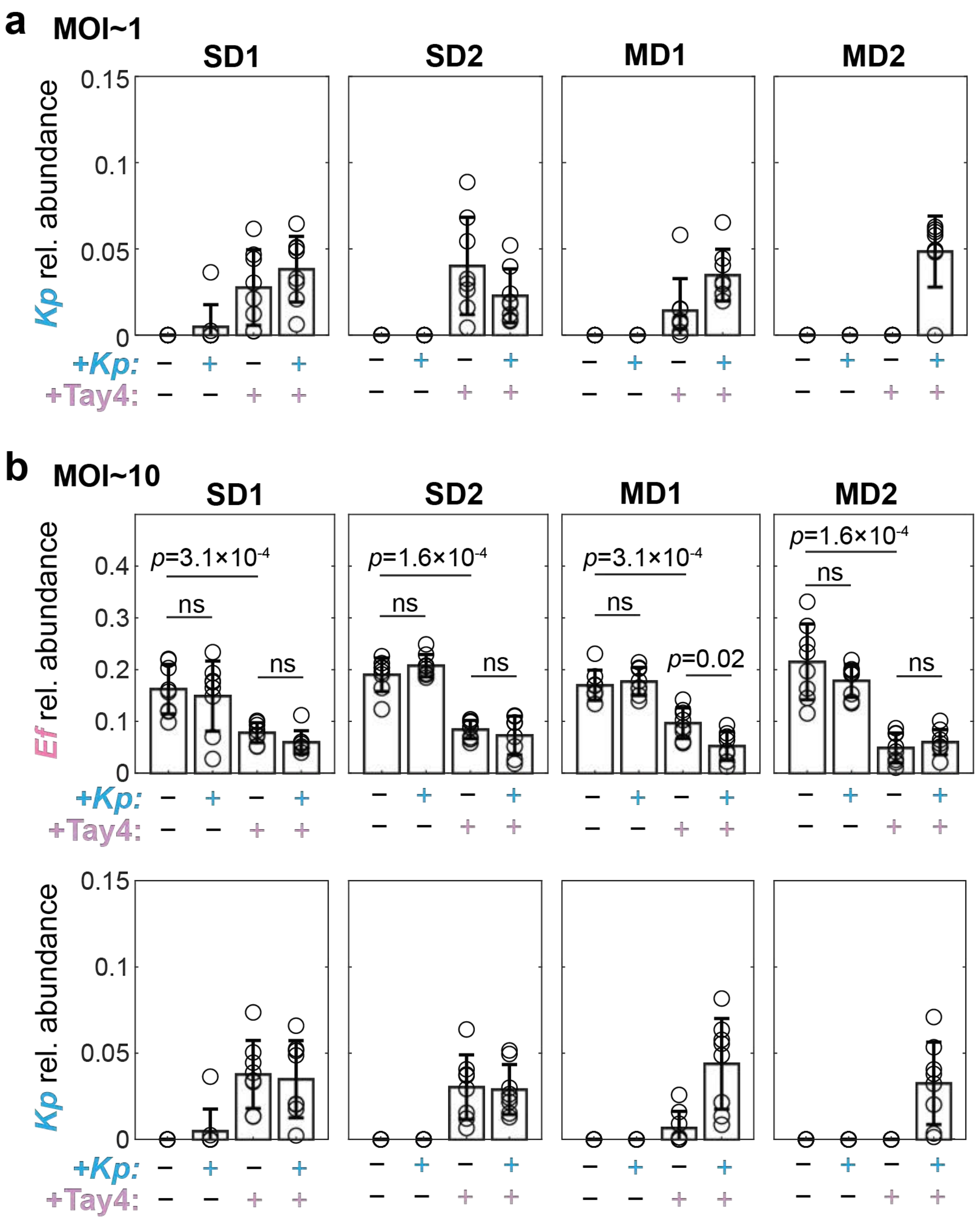


**Figure S6: Addition of *Kp* does not enhance phage-mediated knockout of *Ef*.**

a) Relative abundance of *Kp* in communities from the experiment in **Fig. 3b**. *Kp* abundance increased following Tay4 treatment (MOI~1) across all communities, including MD2, which lacks native *Kp*. Bars show mean±s.d. (*n*=8 biological replicates), with individual replicates overlaid. Addition of *Kp* did not enhance *Ef* knockout.

b) Increasing phage dose does not improve knockout efficacy. Relative abundance of *Ef* (top) and *Kp* (bottom) is shown for communities treated with Tay4 at MOI~10, with or without *Kp* addition. Bars show mean±s.d. (*n*=8 biological replicates) with individual replicates overlaid.

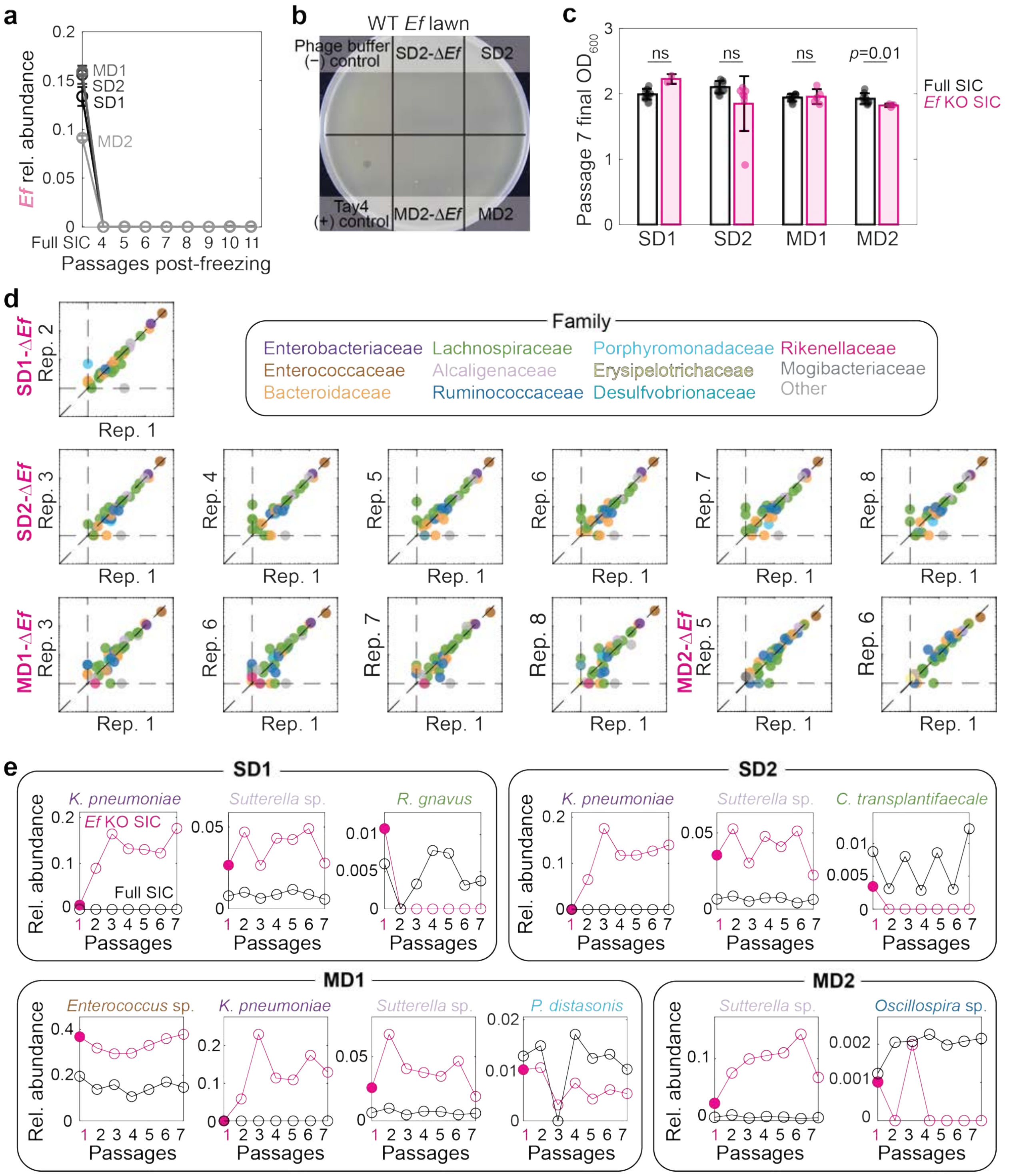


**Figure S7: *Ef* knockout communities are reproducible, stable, and exhibit consistent compositional shifts.**

a) Stability of *Ef* knockout following freezing and passaging. *Ef* relative abundance remained below the limit of detection across multiple passages post-freezing in all Δ*Ef* communities.
b) Absence of residual phage activity in Δ*Ef* communities. Phage lysates derived from Δ*Ef* communities do not form plaques on a wild-type *Ef* lawn, indicating lack of detectable lytic phage.
c) Community biomass following *Ef* knockout. Final $OD_{600}$ at passage 7 is shown for complete SICs (black) and Δ*Ef* communities (pink). No significant differences were observed for SD1, SD2, and MD1, with a modest difference in MD2. Bars show mean±s.d. (*n*=3 biological replicates).
d) Reproducibility of Δ*Ef* community composition across biological replicates. Pairwise comparisons of ASV relative abundance between replicate communities show strong agreement across Δ*Ef* communities. Points represent individual ASVs colored by taxonomic family. Replicate 1 of each community was used for downstream experiments.
e) Consistent taxon-specific responses to *Ef* knockout. Relative abundance trajectories are shown for selected taxa that increase in Δ*Ef* communities (magenta) compared to complete SICs (black). One biological replicate across 7 passages is shown.

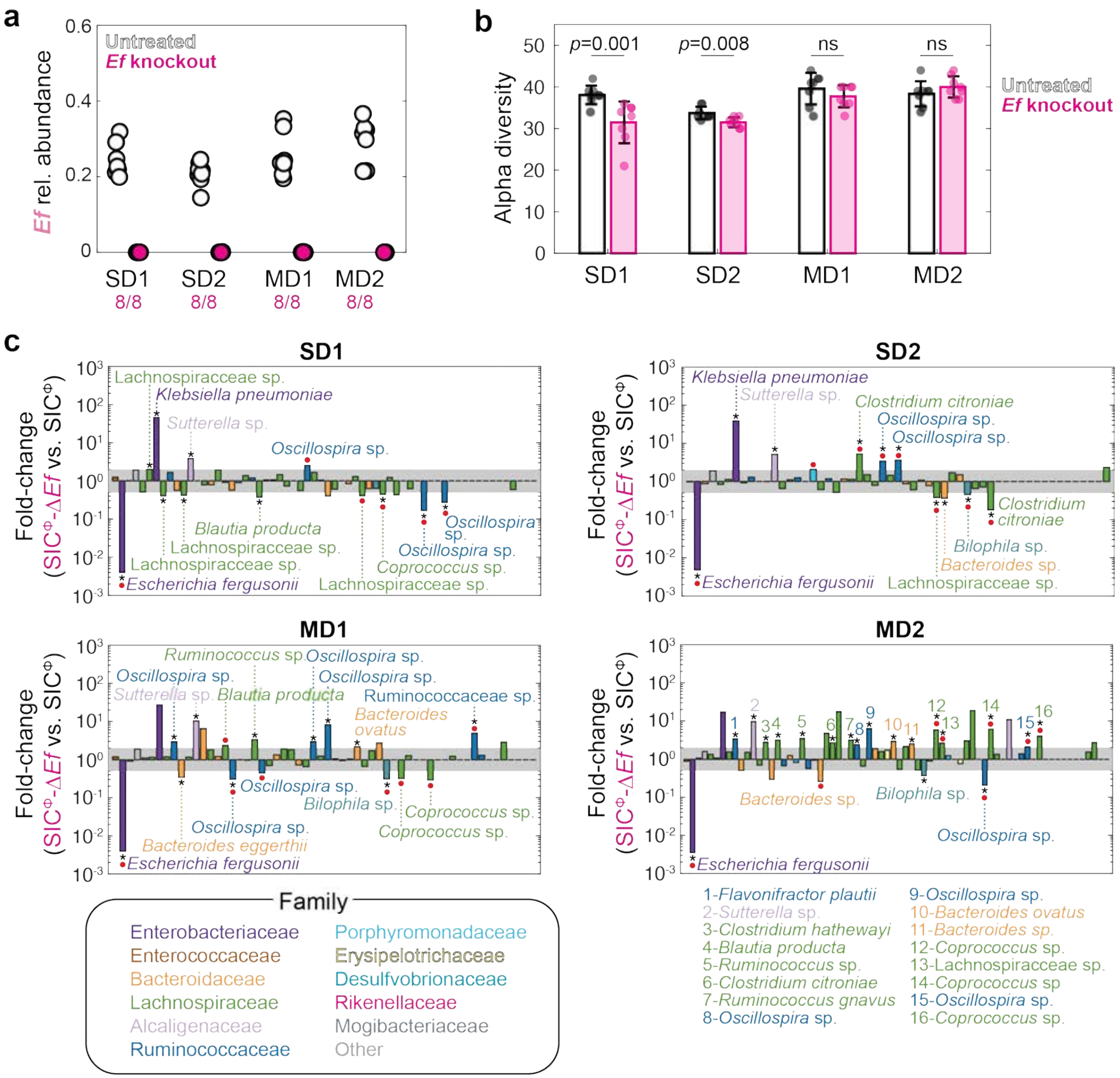

**Figure S8: Phage treatment during community re-derivation enables robust elimination of *Ef* with modest, community-specific effects on diversity and composition.**

a) *Ef* is eliminated when communities are re-derived from stool under phage treatment. Stool samples were treated with a phage cocktail during passage 1 and passaged without phage for six additional passages. *Ef* relative abundance at passage 7 is shown for untreated communities (white) and phage-treated communities (magenta), in which *Ef* is below the limit of detection in all replicates (*n*=8 biological replicates), suggesting that knockout is more efficient when *Ef* starts at low abundance.

b) Alpha diversity following *Ef* removal. Diversity, measured as the number of unique ASVs, is modestly reduced in SD communities and unchanged in MD communities. Bars show mean±s.d. (*n*=8 biological replicates), with individual replicates overlaid. *p*-values were calculated using two-sided Wilcoxon-Mann-Whitney tests (ns: not significant, $p>0.05$).

c) Community composition following re-derivation with phage. Fold change in ASV abundance is shown for Δ*Ef* communities (phage-treated) relative to corresponding untreated communities. Bars represent mean fold change across biological replicates (*n*=8) after passage 7. The shaded region indicates a 2-fold increase/decrease. ASVs with significant differences (two-sided Wilcoxon-Mann-Whitney test with Benjamini-Hochberg correction, adjusted $p<0.05$) are marked with asterisks. ASVs that newly emerge above or drop below the limit of detection ($10^{-3}$) in >4 replicates are highlighted (red circles). Colors denote taxonomic family.

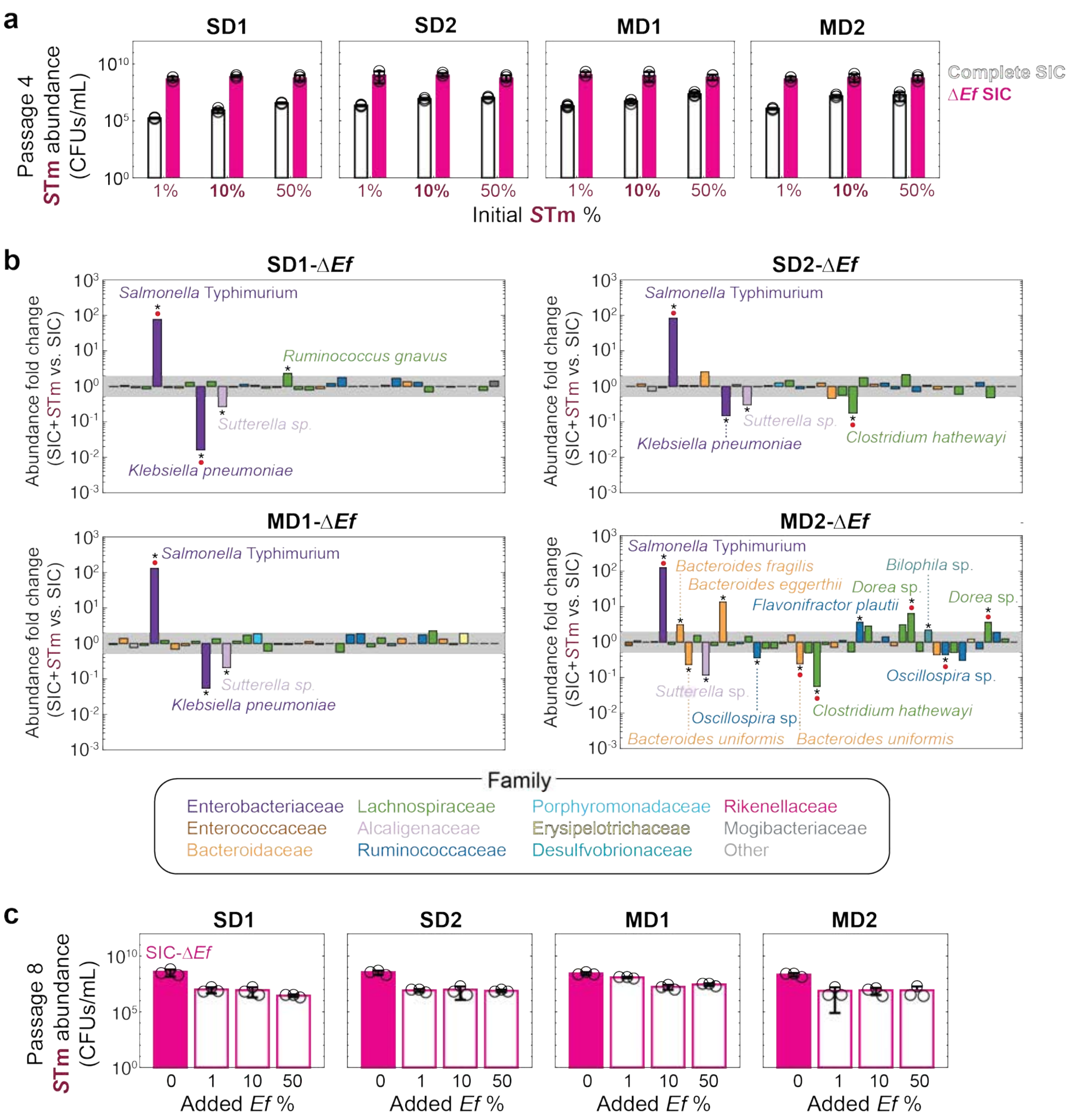


**Figure S9: *Ef* is necessary and sufficient to suppress *Salmonella* invasion across communities.**

a) *S*Tm abundance following invasion of complete and Δ*Ef* communities. Absolute abundance (CFUs/mL) of *S*Tm is shown after four passages for initial inoculation fractions of 1%, 10%, and 50% into complete communities (white bars) or Δ*Ef* communities (magenta). Bars show mean±s.d. (*n*=3 biological replicates), with individual replicates overlaid. The 10% condition (bolded) was for subsequent reintroduction experiments.

b) Community compositional changes following *S*Tm invasion of Δ*Ef* communities. Fold change in ASV abundance is shown for *S*Tm-invaded versus uninvaded communities (10% initial *S*Tm). Bars represent mean across biological replicates (*n*=3) after passage 4. The shaded region represents a 2-fold increase/decrease. ASVs outside this region with significant differences (two-sample Student's *t*-tests with Benjamini-Hochberg correction, adjusted $p<0.05$) are marked with asterisks. ASVs that newly emerge above or drop below the limit of detection ($10^{-3}$) between conditions are highlighted (red circles). Colors denote taxonomic family.

c) Reintroduction of *Ef* suppresses established *S*Tm. *S*Tm abundance (CFUs/mL) is shown after reintroduction of *Ef* into Δ*Ef* communities previously invaded with 10% *S*Tm. *Ef* was added at 1%, 10%, or 50% at passage 5, and communities were passaged to steady state (passage 8). *S*Tm abundance decreased following *Ef* reintroduction across all communities. Bars show mean±s.d. (*n*=3 biological replicates), with individual replicates overlaid.

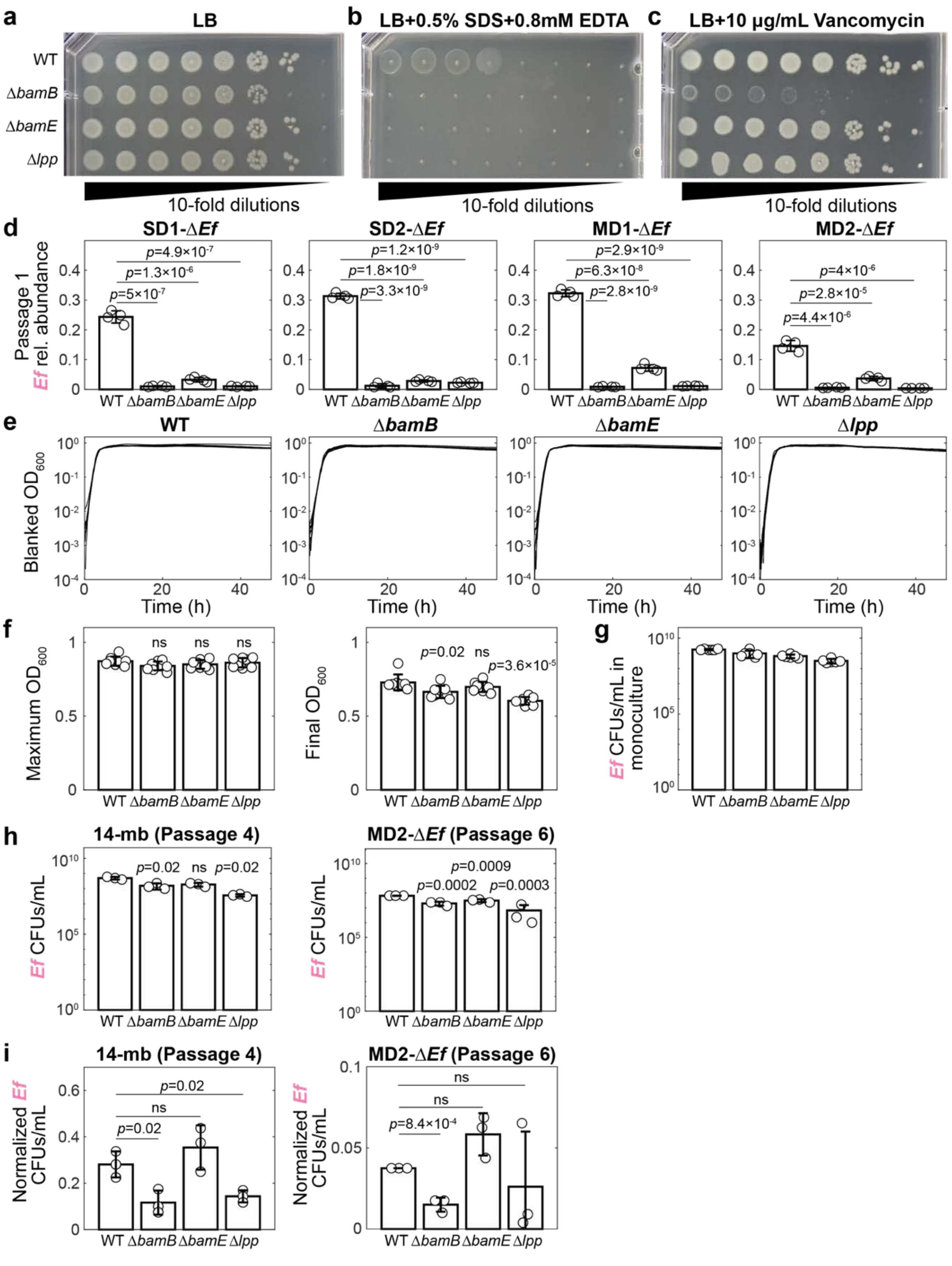


**Figure S10: Outer membrane mutants exhibit increased permeability and reduced fitness specifically in community contexts.**

a-c) Phenotypic characterization of outer membrane mutants. Ten-fold serial dilutions of wild-type (WT) *Ef* and deletion mutants (Δ*bamB*, Δ*bamE*, Δ*lpp*) were plated after 24 h of anaerobic growth. (a) Growth on LB agar shows similar colony formation across strains. (b) Growth on LB supplemented with 0.5% SDS and 0.8 mM EDTA reveals increased membrane permeability in mutants. (c) Growth on LB supplemented with 10 μg/mL vancomycin shows increased sensitivity for Δ*bamB*.

d) Early fitness defect of outer membrane mutants in communities. Relative abundance of wild-type and mutant *Ef* strains following co-culture with Δ*Ef* communities after passage 1. Bars show mean±s.d. (*n*=4 biological replicates), with individual replicates overlaid. *p*-values were calculated using two-sample Student's *t*-tests.

e-g) Growth of outer membrane mutants in monoculture. Bars represent mean±s.d. (*n*=8 biological replicates), with individual replicates overlaid. *p*-values were calculated using two-sample Student's *t*-tests. (e) Growth curves in BHI show similar dynamics across strains. Curves are biological replicates (*n*=8). (f) Maximum and final $OD_{600}$ (background-subtracted) are largely comparable, with minor reductions in some mutants. (g) CFU measurements after 48 h confirm similar overall growth yields (*n*=6 biological replicates). Together, these data indicate minimal intrinsic fitness defects in monoculture.

a) Reduced abundance of outer membrane mutants during co-culture. CFUs/mL are shown after co-culture with a 14-member (14-mb) synthetic community (left; passage 4) or MD2-Δ*Ef* community (right; passage 6). Bars show mean±s.d. (*n*=3 biological replicates), with individual replicates overlaid. *p*-values were calculated using two-sample Student's *t*-test between WT and each mutant.

b) Community-specific fitness defects of outer membrane mutants. CFUs/mL from (**h**) normalized to monoculture values reveal reduced relative fitness in community contexts. Bars show mean±s.d. (*n*=3 biological replicates), with individual replicates overlaid. *p*-values were calculated using two-sample Student's *t*-tests.

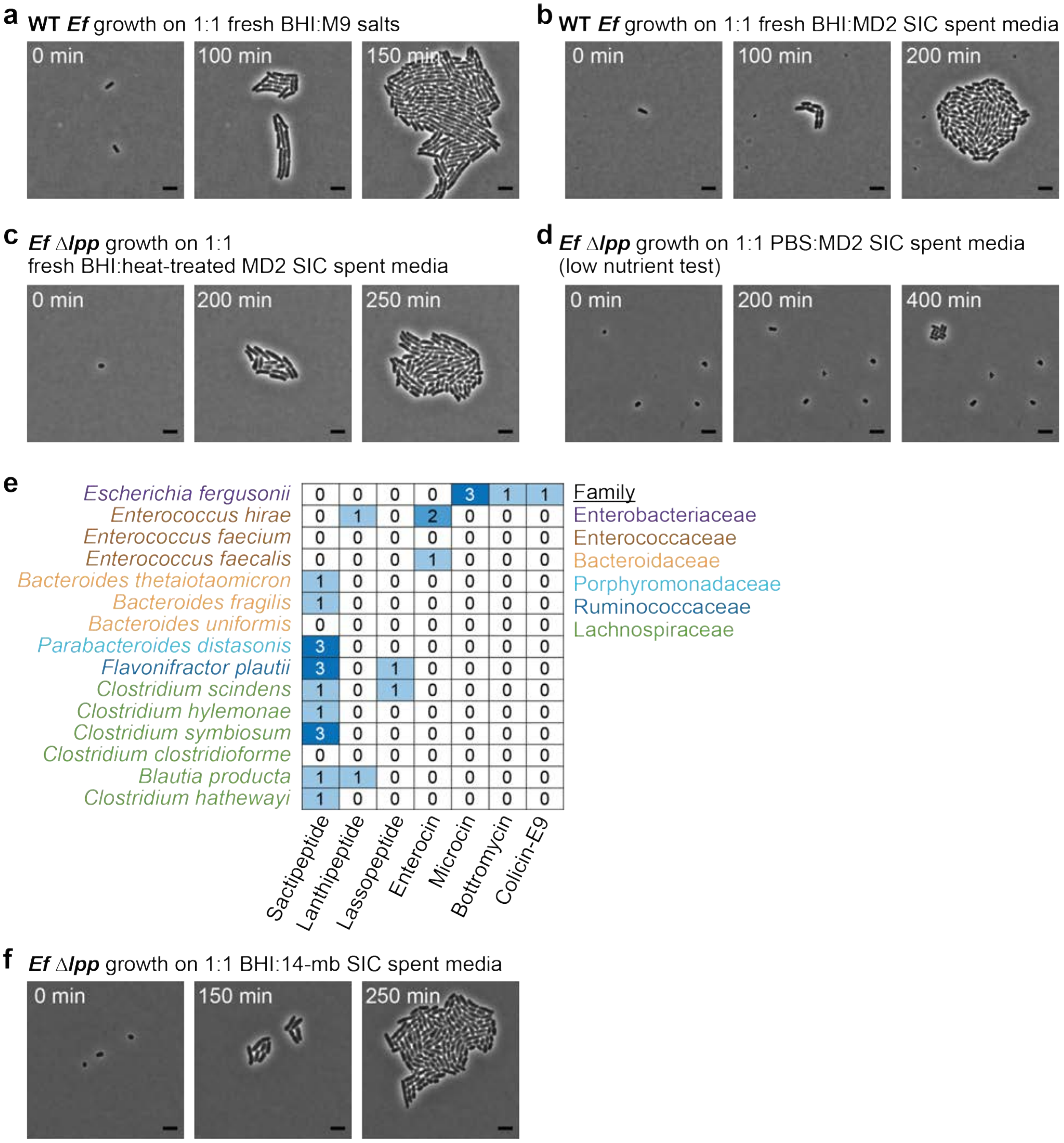


**Figure S11: The Δ*lpp* growth defect depends on heat-labile factors in MD2 spent medium and is not recapitulated by nutrient limitation or defined communities.**

a) Time-lapse microscopy of wild-type (WT) *Ef* grown on 1% agarose pads containing 50% BHI and 50% M9 salts (**Movie S3**). Cells exhibit normal growth and morphology. Scale bar: 5 µm.

b) Wild-type *Ef* growth on 1% agarose pads containing 50% fresh BHI and 50% MD2-Δ*Ef* spent medium (**Movie S5**). Cells maintained normal morphology, indicating that community spent medium does not perturb wild-type growth. Scale bar: 5 μm.

c) *Ef* Δ*lpp* growth on 1% agarose pads containing 50% fresh BHI and 50% heat-treated MD2-Δ*Ef* spent medium (**Movie S6**). Heat treatment (95 °C, 15 min) abolished the morphological defects observed in untreated spent medium, indicating that the active factor is heat-labile. Scale bar: 5 μm.

d) *Ef* Δ*lpp* growth on 1% agarose pads containing 50% PBS and 50% MD2-Δ*Ef* spent medium (**Movie S7**). Under low-nutrient conditions, Δ*lpp* cells showed minimal growth without morphological defects. Scale bar: 5 μm.

e) Predicted ribosomally synthesized and post-translationally modified peptides (RiPPs) encoded by members of the 15-species community, identified using BAGEL4[83].

f) *Ef* Δ*lpp* growth on 1% agarose pads containing 50% BHI and 50% spent medium from the 14-member (14-mb) synthetic community (**Movie S7**). Unlike MD2-Δ*Ef* spent medium, 14-member spent medium did not induce morphological defects, despite reduced fitness in co-culture (**Fig. 6g**). Scale bar: 5 μm.
